# Modulation of biphasic pattern of cortical reorganization in spinal cord transected rats by external magnetic fields

**DOI:** 10.1101/2024.04.10.588799

**Authors:** Sajeev Kaur, Kanwal Preet Kochhar, Suman Jain

**Affiliations:** Department of Physiology, All India Institute of Medical Sciences, Ansari Nagar East, New Delhi-110029, India

**Author notes:** **Corresponding Author:** Dr. Suman Jain Professor, Department of Physiology, All India Institute of Medical Sciences, Ansari NagarEast, New Delhi-110029, India.

**Keywords:** spinal cord injury, cortical reorganization, plasticity, brain derived neurotrophic factor (BDNF), extremely low frequency magnetic field (ELF-MF), Nogo-A, grip strength, allodynia

## Abstract

Magnetic field induces electric field in the brain and has the potential to modulate cortical plasticity giving rise to sustained excitability as well as behavioral modifications. The present study demonstrates the effect of extremely low frequency magnetic field (ELF-MF) on cortical reorganization after SCI in complete transection rat model. The spinal cord of male albino Wistar rats was completed transected at T-13 spinal level and ELF-MF (50 Hz, 17.96 µT) exposure given either for 5 or 12 or 32 days for temporal observations. A significant functional recovery was evident in locomotor, sensorimotor and motor behavioral tests after 32 days of magnetic field exposure. This was associated with an amelioration in cortical electrical activity, decrease in the lesion area and volume and modulation of plasticity associated proteins like Nogo-A and BDNF in a biphasic pattern. The results suggest improvement of functional, electrical and morphological parameters along with upregulation of plasticity associated proteins in an in-vivo SCI rat model following ELF-MF exposure for 32 days but not after 5, 12 days exposure

## Introduction

Cortical maps undergo reorganization in response to various internal as well external stimuli, e.g. use or disuse, skill learning, and injuries like Spinal cord injury (SCI) signifying capacity of adult brain for substantial plasticity (Pons et al., 1991; Jain et al., 1997; Florence et al., 1998; Endo et al., 2007; Ghosh et al., 2009; Holtmaat and Svoboda, 2009; Barnes and Finnerty, 2010;). In SCI an alteration in cortical plasticity is mainly due to irreversible as well as abrupt deafferentation of cortical circuitry both immediately as well as chronically (Aguilar et al., 2010). Within 3 days of SCI, an invasion of cortical representation of the forelimb area in somatosensory cortex was observed in adjacent sensory-deafferented hindlimb area (Endo et al., 2007) The BOLD (blood oxygen level dependent) signals evoked during fMRI studies while stimulating forelimb demonstrated striking biphasic responses in the somatosensory cortex of SCI rats, an increase at 3 days followed by temporary decrease at 1-2 weeks and at 1-6 months permanent increase, suggesting reorganization of forelimb cortical area after SCI (Endo et al., 2007). The mechanism of reorganization of cortical inputs has been tracked back at subcortical levels especially, thalamus and brain stem (Tandon et al., 2008; Dutta et al., 2014; Halder et al., 2018). Whilst the cortical plasticity is important for functional recovery but massive or aberrant reorganization can cause pathological conditions such as neuropathic pain and phantom sensations (Yague et al., 2014). Thus, understanding the basis of cortical plasticity is imperative to maximize good over bad side to optimize cost effective therapies that maximize functional recovery and minimize neuropathic pain following SCI (Moxon et al., 2014). There are several factors that determine the reorganization of the cortex, for eg; animal species, time elapsed after the injury, age of the animal or subject, and intervention given after the injury. The process is also accompanied with an alteration of plasticity associated proteins like Nogo-A and BDNF which forms the molecular basis of such reorganization/ plasticity. Nogo-A is a membrane bound protein that is recognized as an inhibitor of CNS specific axonal regeneration. Current studies have deciphered the modulatory role of Nogo proteins and their receptors in the neurite growth and branching in the developing brain as well as growth-restricting function during maturation of CNS. Thus, the function of Nogo in CNS is now understood to be that of a negative regulator of neuronal growth, leading to stabilization of the CNS wiring at the expense of extensive plastic rearrangements and regeneration after injury (Schweigreiter & Bandtlow, 2006; Wang et al., 2015). BDNF is a member of the neurotrophin family which is imperative for the growth and survival of neurons, modulation of neurotransmitters, neurogenesis, and synaptic plasticity. Thus, BDNF plays a pivotal role in development, brain plasticity, and is indispensable for learning and memory (Mattson et al., 2004; Bathina & Das., 2015). The biphasic BOLD responses during cortical reorganization parallel the changes of NgR, LINGO-1, and BDNF gene activity. Endo et al., (2007) found that NgR, its co-receptor LINGO-1, which is an inhibitory protein is down-regulated especially in cortical areas deprived of sensory input and in adjacent cortex immediately after injury while BDNF is upregulated. Thus, there is an involvement of Nogo signaling in cortical activity-dependent plasticity in the somatosensory system.

Neuromodulatory effects of magnetic fields by inducing eddy currents in the neural tissue have been of immense potential in treatment of various neurological disorders. In hemisection or complete transection SCI rat model, an improvement in sensorimotor and locomotor function (Das et al., 2012), feeding behavior, body weight (Kumar et al., 2010), attenuation of SCI-induced osteoporosis (Manjhi et al., 2013) and tonic pain, restoration of neurotransmitter milieu (Kumar et al., 2013) has been reported after ELF-MF exposure. Electromagnetic field has shown to alter cortical excitability and instigate plastic changes in the brain (Belci et al., 2004). Studies have been done to assess the function of NIBS in SCI patients to maximize the functional outcome (Belci et al., 2004) and in reducing neuropathic pain (Fregni et al., 2006). The low intensity repetitive transcranial magnetic stimulation (LI-rTMS) has shown to promote cortical reorganization in rat model with visual system defects (Rodger et al., 2012; Makowieki et al., 2014). Dileone et al., (2018) studied the long-term effects of transcranial static magnetic field stimulation (tSMS) on the excitability of the motor cortex. However, the effect of ELF-MF (19.46µT, 50Hz) on cortical plasticity at different time points after complete spinal cord injury has not been studied.

In order to gain insight into the effect of ELF-MF (50Hz, 17.96µT) intervention on cortical plasticity after spinal cord injury, we designed a temporal study in complete transection SCI model. The study focuses primarily on two factors; time elapsed after the injury, and effect of extremely low frequency magnetic field (ELF-MF) intervention. We hypothesized that ELF-MF exposure will promote plasticity by modulating expression of Nogo-A and BDNF after SCI leading to functional and electrical recovery.

## Materials and Methods

### Animals

Adult male Wistar rats (250-280g; n= 72) were acquired from the Central Animal Facility (All India Institute of Medical Sciences, New Delhi, India) and kept individually in the polypropylene cages (50cmX 20cmX 15cm) provided with standard husk bedding. Animals were housed at an ambient room temperature of 24±2 °C, relative humidity of 50-55%, light-dark cycle of 14:10 hours, and given laboratory food pellets and clean drinking water *ad libitum.* The ethical approval for the study was obtained from the All India Institute of Medical Sciences, New Delhi, India (884/IAEC/15). Rats were randomly divided into four groups: control (no surgery; n= 18), sham (laminectomy without complete transection; n= 18), SCI (complete transection of spinal cord; n= 18) and SCI+ MF (complete transection along with magnetic field intervention; n= 18). Each group was further divided into three temporal subgroups of 5 days (n= 6), 12 days (n= 6), and 32 days (n= 6), thus making it to a total of 18 rats in each group.

### Complete transection surgery

The rats were given pre-anesthetic glycopyrrolate (20-30 mg/kg; intramuscularly) 10 min prior, followed by anesthesia with pentobarbitone (50 mg/kg; I.P.). The rats were fixed in a stereotaxic instrument (Model-1404, David Kopf, USA), and dorsal surface overlying the thoraco-lumbar vertebra was shaved to expose the skin. A mid-dorsal skin incision was made from the upper thoracic region to the upper lumbar region. The fascia and muscle were removed to expose spinal cord followed by T10-T12 laminectomy. The spinal cord was cut completely at T13 spinal level (T11 vertebrae) using microscissors, and completeness of the incision was ensured using a glass seeker. Absorbable gelatin (AbGel) sponge was kept in the transected region. The muscle layer and skin were sutured (Ethicon, non-absorbable surgical suture, Johnson & Johnson Ltd., Mumbai, India) and neosporin powder was applied to the sutured skin surface. Sterile ringer lactate solution (5-10 ml; S.C.) was administered to prevent dehydration. The rats were kept on a thermostat pad (Harvard/CMA), and the temperature was monitored continuously during surgery. Broad-spectrum antibiotics like gentamicin (30 mg/kg; I.M.) were given to prevent general infection and enrofloxacin (20 mg/kg; I.M.) to prevent urinary tract infection for 5 days. The urinary bladder was manually evacuated three times every day.

### Extremely low frequency magnetic field (ELF-MF) exposure protocol

The magnetic field (17.96µT, 50Hz, 2hr/ day) exposure was started 24 hours after the injury in SCI+MF group for either 5, 12, or 32 days in a indigenously designed magnetic field (MF) chamber (All India Institute of Medical Sciences, New Delhi, India) as described previously (Manjhi et al., 2013).

### Assessment of Urinary Bladder Function

The urinary bladder function scoring was done every day according to the method described by Martin Schwab’s lab (Liebscher et al., 2005) with modifications. The scoring to assess urinary bladder function (UBF) is as follows: spontaneous evacuation= 1, partial assistance for evacuation= 2, total assistance for evacuation= 3, and bursting of bladder= 4. The rats were manually evacuated thrice a day till the recovery of the bladder.

### Assessment of behavioral functions

All the behavioral parameters including BBB scoring, grip strength and von Frey test were performed before the surgery (baseline) and also on day 3, 10, and 30 in subgroups day 5, 12 and 32 respectively.

### Basso Beattie Bresnahan (BBB) scoring

Motor function was assessed using a modified 0-21 points open field locomotor scale, wherein the rat was left in the open field, and its movements were video recorded using a camera for 4 min. The movement of three joints (knee, ankle, and hip joint), paw placement, weight support, and forelimb/ hindlimb coordination were observed. The movements were scored and analyzed manually by methodology established by Basso, Beattie, and Bresnahan (Basso et al., 1995).

### Grip Strength

The grip strength test was used to assess the muscular strength of the forelimb by recording the amount of force (g) exerted before the release of the grasp from the metallic T-bar of the grip strength meter (IITC Life Science Inc., Woodland Hills, CA). Grip strength procedure consisted of two phases: training and testing. Five training trials were given for two days, followed by testing. Five readings were taken in each trial with an interval of 2 minutes, the median and the two nearest values were averaged (Sandrow-Feinberg et al., 2009).

### von Frey

The purpose of the test was to determine the tactile allodynia below and above the level of lesion. The von Frey test involves two phases: habituation/acclimatization phase and testing phase. The rat was acclimatized in the testing room for 15min and given palatable food (black gram) to eat, followed by habituation in the partial restrainer for 15 min. The habituation was done for three days followed by testing on the fourth day. One the testing day, the instrument was calibrated with the test weight (5.2g) provided by company (IITC Life Science). The rat was habituated as mentioned in acclimatization phase, followed by application of the test stimulus for maximum of 5 sec between the footpads of the rat paw using a rigid tip (constant weight). If no response occurs within 5 sec then the stimulation was done again after 30s. A positive response includes the sudden withdrawal of limb, flinching, licking, or scratching the area where the tip was touched. The withdrawal thresholds (gm) were recorded, and the median value and the two nearest values were averaged. Touching of any other part of the footpads was avoided as the skin’s sensitivity is much lower in those areas (Leem et al., 2012). Five positive trials were noted for each paw with an interstimulus interval of 30sec between each trial to prevent the adaptation of sensory receptors.

### Assessment of cortical electrical activity (EEG)

The cortical electrical activity was recorded on day 5, 12, and 32 (terminal days) in subgroups day 5, 12 and 32 respectively. The rat was anesthetized using urethane (1.5 g/kg; I.P.), fixed in the stereotaxic apparatus (model 1404, David Kopf, Tujunga, CA, USA) and scalp, cranium Burr holes were made and screw electrodes implanted in frontal (AP +2, ML 3.5) as well in parietal (AP -2, ML 3.5) cortex in both the hemispheres using bregma as reference. The reference electrode was placed in the frontal lobe and parietal lobe respectively near the midline. The size of the watch screw was chosen so as to ensure that dura does not get compressed or damaged while inserting electrodes. Filters were applied to reduce noise (Band pass 0.5-35Hz, Notch filter: 50 Hz). EEG recording was done at sampling rate of 400S/s for 10 min using LabChart 8 software (AD Instruments, USA).Offline digital filters were applied after acquisition for delta (θ), theta (θ), alpha (α), and beta (β) to separate the different components as per their band frequency (Sitnikova et al., 2005). The total power for each wave was calculated using the same software.

### Histological assessment

The rats were sacrificed by transcardial perfusion under deep anesthesia using normal saline followed by 4% paraformaldehyde (PFA). The spinal cord along with vertebral column (T5-L5) and brain were isolated at the end of the study period at day 5, 12 and 32 in the respective subgroups. Both the tissues were post fixed in PFA for 24 h at 4°C.

### Lesion area and volume

Lesion area and volume was calculated to determine the extent of lesion in the spinal cord. Spinal cord was cryoprotected in 15% sucrose for 24 h and 30% sucrose for 24-48 hours serially. The alternate longitudinal sections of 16µm were taken on poly-l-lysine coated slides using cryotome. Cresyl violet staining was done using standard protocol to identify the basic neuronal structures and lesion area in the spinal cord. The stained sections were visualized under bright microscope (Nikon Upright Motorized microscope ECLIPSE Ni-E, Japan).

To estimate the extent of lesion in the spinal cord, three slides with four sections on it were taken. In each cresyl violet stained section, lesion area (mm^2^) was traced (Nikon Upright Motorized microscope ECLIPSE Ni-E, Japan) by selecting the affected area using NIS-Element software. To calculate the total volume of the lesion, the total averaged area was multiplied with the number of sections per spinal cord and the thickness of single section. Lesion volume (mm^3^) = Lesion area X 16 (thickness of section in µm) X12 (total sections) X4 (sections on each slide) X2 (factor to indicate that every alternate section was taken).

### Immunofluorescence and signal counting

6mm block of brain was embedded in cryomatrix and every 5^th^ coronal section of 16µm was taken. Immunofluorescence was done on brain sections to assess expression of plasticity associated proteins Nogo-A and BDNF with rabbit anti-Nogo-A (1:200; Invitrogen, USA), rabbit anti-BDNF (1:500; Invitrogen,USA), and anti-rabbit FITC labeled 2° antibody (1:400; Vectors Laboratory, USA).

Four antibody stained sections were taken for each rat for analysis depending on the coordinates containing M1 (primary motor cortex), M2 (secondary motor cortex), S1Fl (somatosensory, forelimb area) S1HL (Somatosensory, hindlimb area) area from the rat’s atlas (Paxinos and Watson atlas, sixth edition 2007) (Figure 1). Imaging of all the areas was done in two different fields in all four sections using NIS-Element software (Nikon Upright Motorized microscope ECLIPSE Ni-E, Japan) (Endo et al., 2007). Image J software was used for counting of the signals and Nogo-A+ and BDNF+ cells per field were noted. Average was taken for cell counts/ field for both the fields for each area for all four sections.

**Figure 1:**
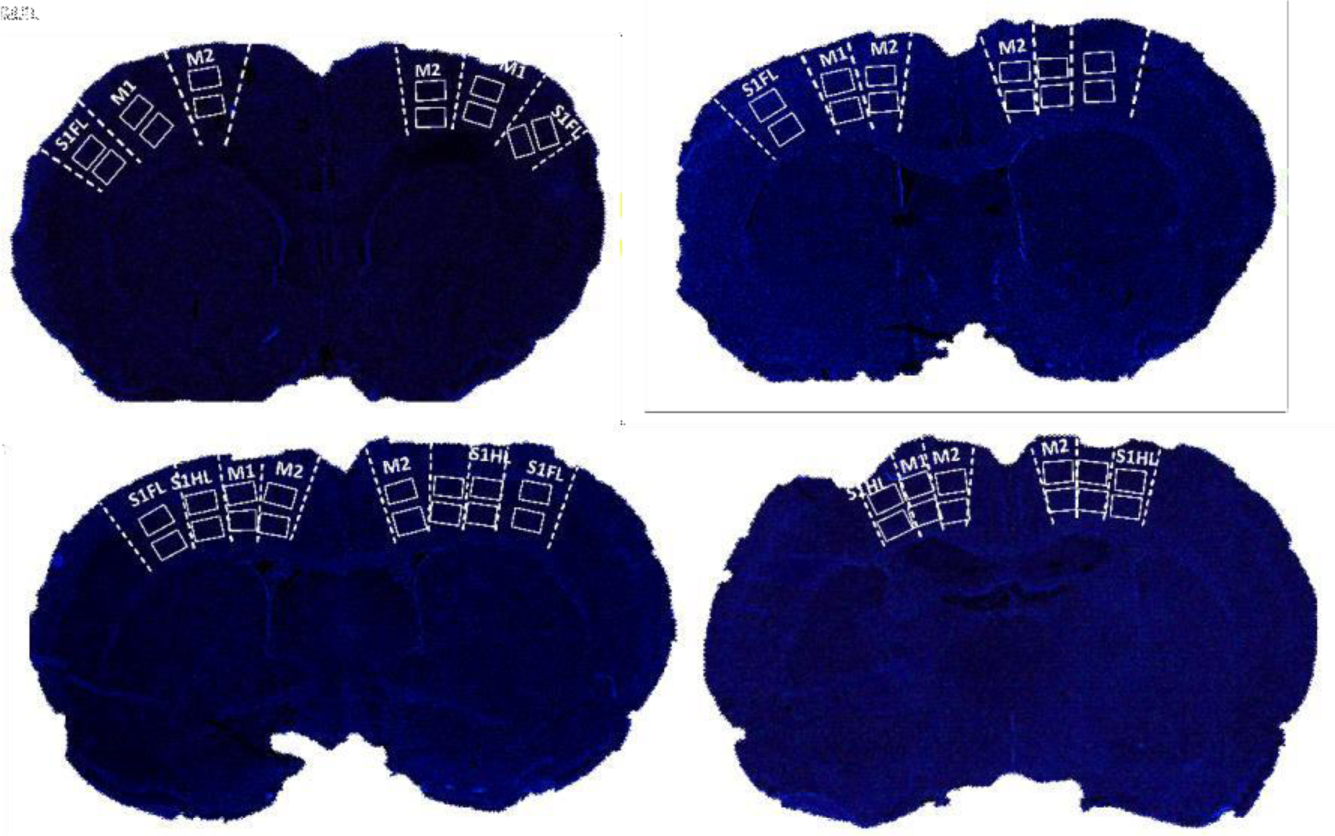
Areas localization procedure using NIS software for Nogo-A and BDNF. White boxes showed fields (top, field 1; bottom, field 2). M1= primary motor cortex; M2= secondary motor cortex; S1Fl= Somatosensory, forelimb area; S1HL= Somatosensory, hindlimb area.

### Statistical analysis

Statistical analysis was done using GraphPad Prism 5. For the intergroup analysis, one way ANOVA (nonparametric: no assumption of Gaussian distribution) - Kruskal-Wallis test and post hoc analysis was done by Dunns (all groups included); significance: 0.05 (95% confidence interval).The t-test (non-parametric) for comparison between post op-SCI and SCI+MF group: Mann Whitney test. P values: Two-tailed (95% confidence interval). For the intragroup analysis, 2 Way ANOVA. Tukey’s Multiple comparison test.

## Results

### Assessment of autonomic function

Urinary bladder function scoring was done to assess the effect of SCI on the autonomic function. All the animals recruited for the study were able to evacuate the urine spontaneously before the injury, thus UBF score was 1. After the complete transection injury, there was a complete loss of spontaneous evacuation of urine (UBF score= 3) in SCI and SCI+ MF groups. In both these groups, complete recovery of bladder function did not occur till the end of the study period (32 days), but partial recovery (UBF score =2) was evident around 24-29 days in SCI and 16-20 days in the SCI+MF group. The recovery of the bladder was significantly higher (p=0.0050) in the MF group in comparison to the SCI group (Figure 2).

**Figure 2:**
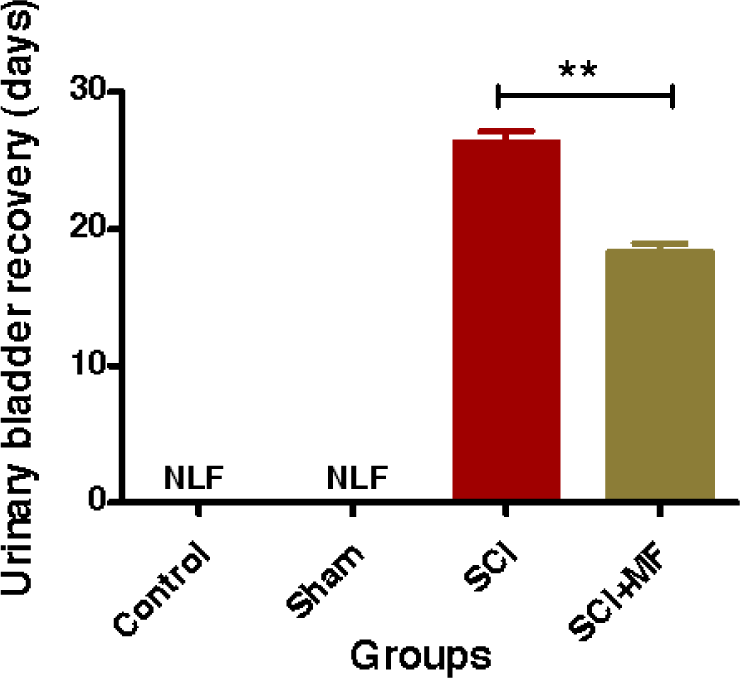
Intergroup comparison of urinary bladder function after MF exposure in 32 days subgroup. Data expressed as mean± SEM (n=6/ group). NLF= No loss of function. * indicates comparison between SCI and SCI+MF groups; p≤ 0.01.

### Assessment of behavioral parameters

To assess the behavioral parameters three different types of comparisons were done: intragroup comparison (baseline vs respective days), intragroup temporal comparison, and intergroup comparison.

### Locomotor function

The pre-surgery BBB score was 21 for all the rats which indicated normal locomotor function. After surgery, rats exhibited complete paraplegia and moved using forelimbs whilst their hindlimbs were dragging against the floor of their home cage. In both SCI and SCI+MF groups, but not in sham group, there was a decrease in the BBB score to zero right after the surgery leading to no perceptible hindlimb movements. There was a gradual improvement in BBB score from day 3 to 30 after surgery in both SCI and SCI+MF, suggestive of minimal movements in the hindlimb joints. Intragroup comparison (with baseline) showed significant decrease in BBB score on day 3 (p≤ 0.001), 10 (p≤ 0.001) and 30 (p≤ 0.001) in SCI group when compared to baseline and the pattern was same in SCI+MF group as well at day 3 (p≤ 0.01), 10 (p≤ 0.05), and 30 (p≤ 0.05). Intragroup temporal comparison was done using two-way ANOVA. The time point (row factor; p<0.0001; F=63; df= 2) and groups (column factor; p<0.0001 F=15; df=3) and the interaction between the two factors (p<0.0001 F=21; df=6) found significant decrease in BBB score after injury which improved in both the groups over the period of the time. Intergroup comparison revealed significant effect of MF intervention, BBB score was higher in SCI+MF group as compared to SCI group on day 10 (p= 0.0043) and day 30 (p= 0.0153) (Figure 3).

**Figure 3:**
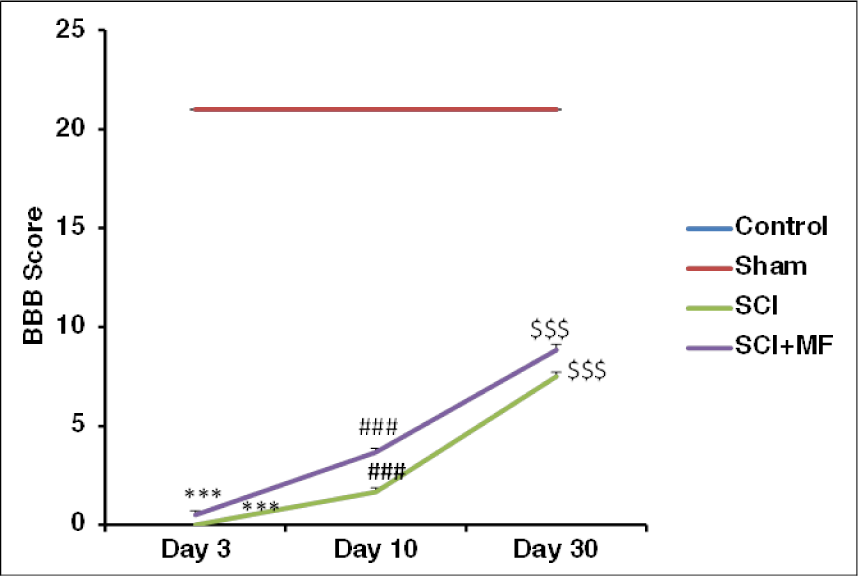
(a) Intragroup temporal comparison of BBB locomotor function score; * (b) Intergroup comparison of BBB scores at different time points after MF exposure. Day 3 (A); day 10 (B); day 30 (C). Data expressed as Mean±SEM (n=6/ group). Day 3-Day 10; ^#^Day 10-Day 30; ^$^Day 3-Day 30. ^*/#/$^ ≤0.05; ^**/##/$$^ ≤0.01, ^***/###/$$$^ ≤0.001.

### Assessment of muscular strength

In both SCI and SCI+MF groups, paraplegia occurred after surgery and thus, animals were moving in the cages using their forelimbs. Intragroup comparison (with baseline) showed significant increase in forelimb muscular strength in SCI group at day 10 (p≤ 0.01) and day 30 (p≤ 0.05). In two way ANOVA done for intragroup temporal comparison the time point (row factor; p< 0.0001; F=39; df=3) and force in grams (column factor; p<0.0001; F=22; df=2) and the interaction between these two factors (p<0.0001; F=20; df=6) showed significant increase in the muscular strength in SCI group over the course of the study showing the effect of SCI on the forearm muscular strength but not in SCI+ MF group. Intragroup temporal comparison in SCI, showed a significant increase in muscular strength from 402.28 ± 9.48 at day 3 to 574.53 ±13.01 at day 30 (p ≤ 0.001), while in SCI+MF group there was no significant increase in grip strength from 389.97±11.83 at day 3 to 418.00± 6.99 at day 30.

Intergroup comparison showed significant increase in muscular strength in SCI group in comparison to control and sham group on day 10 and day 30 but there was no significant difference between SCI+MF in comparison to control and sham on any of the days. Intergroup comparison (t-test) between SCI and SCI+MF showed that muscular strength (force in g) was significantly decreased in SCI+MF group as compared to SCI group on day 10 (p=0.015) and day 30(p=0.002), suggesting recovery (Figure 4).

**Figure 4:**
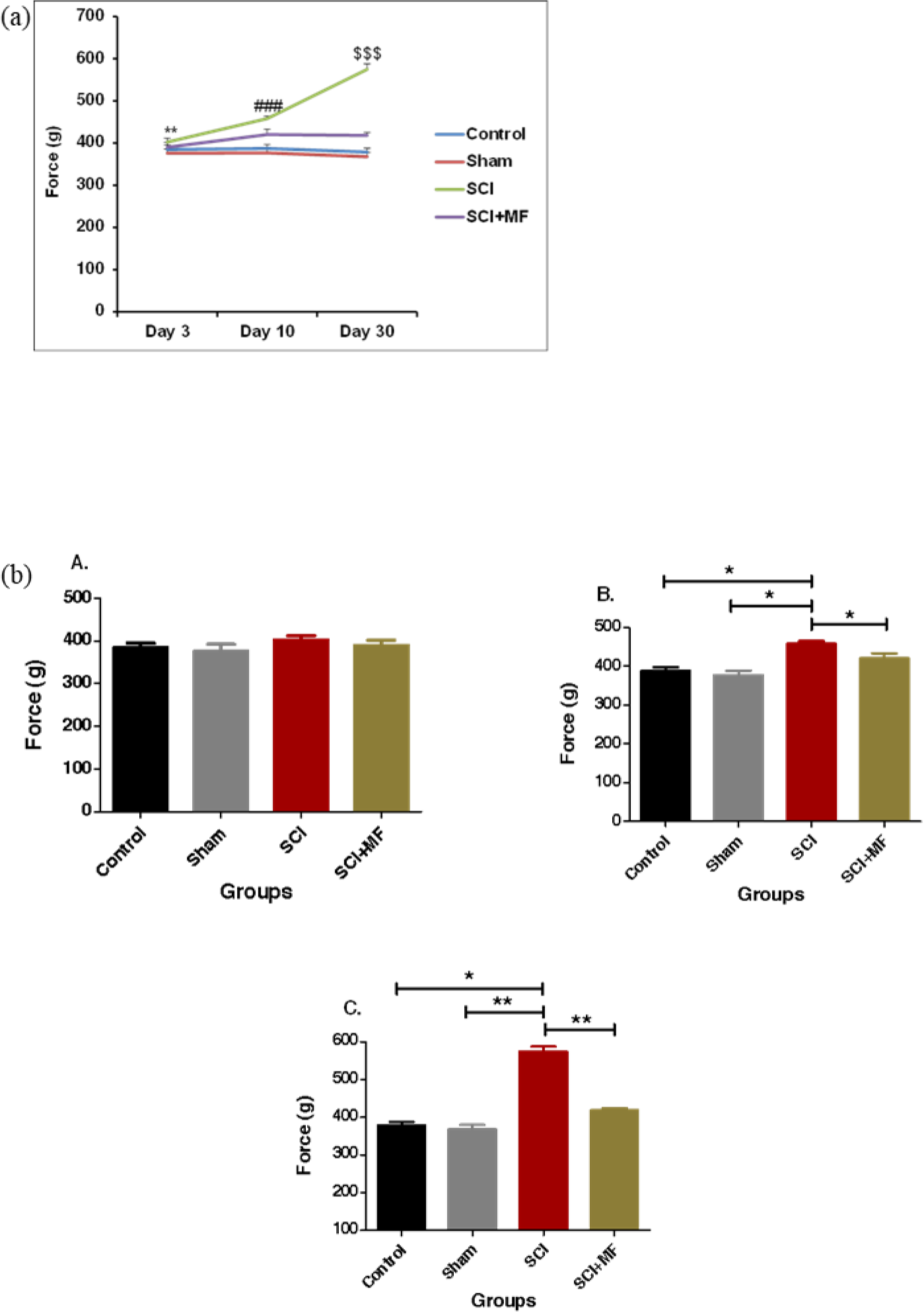
(a) Intragroup temporal comparison of forelimb muscular strength (force (g); (b) Intergroup comparison of forelimb muscular strength expressed as force (g) at different time points after MF exposure. Day3 (A); Day10 (B); Day30 (C). * indicates comparison between day 3 and day 10; ^#^ indicates comparison between day 10 and day 30; ^$^ indicates comparison between day 3 and day 30.Data expressed as Mean±SEM (n=6/ group). ^*/#/$^ ≤0.05; ^**/##/$$^ ≤0.01, ^***/###/$$$^ ≤0.001.

### Assessment of tactile allodynia

There was no significant difference between the baseline values among the groups (baseline values ranged from 10-21 g for forepaws and 24-39 g for hindpaws). Animals showing any kind of aberrant behaviors during baseline recording were removed from the study. von Frey test could not be done (CD) in right and left hindpaw (RH; LH) on day 3 and 10 because there was no plantar placement of the hindpaws in these animals. Intragroup comparison (with baseline) showed increase in withdrawal threshold in SCI group from baseline at day 3 (RF; p ≤ 0.01, LF; p ≤ 0.05), day 10 (RF; p ≤ 0.05, LF; p ≤ 0.05), and day 32 (RF; p ≤ 0.05,LF; p ≤ 0.05) in forepaws, whereas, there was decrease in threshold in SCI group on day 30 (RH; p ≤ 0.05, LH; p ≤ 0.05) for hindpaws.

Intragroup temporal comparison was done to study the changes in tactile sensations in all the groups using two-way ANOVA. For right forepaw and left forepaw, the time point (row factor; RF; P = 0.8408, F=0.174, df=2; LF; p=0.065,F=2.93, df=2) and withdrawal threshold in grams (column factor; RF; p< 0.0001, F=69, df=3; LF; p< 0.0001, F=78.45, df=3), and the interaction between these two factors (RF; p=.0002, F= 5.73, df=6; LF; p< 0.0003; F= 5.44; df=6) showed significant increase in withdrawal threshold after the SCI but decrease in withdrawal threshold in SCI+MF group. For hindpaws, intragroup temporal comparison is not shown because the test was not performed on day 3 and 10. For forepaws, intragroup temporal comparison in SCI, showed a significant increase in withdrawal threshold (p ≤ 0.001; p≤ 0.001) in right forepaw and left forepaw from day 3 (RF; 24.20±1.02, LF; 24.01± 1.08) to day 30 (RF; 30.03±0.12, LF; 30.05±0.67) while in SCI+MF group there was significant (p ≤ 0.01; p≤ 0.01) decrease in withdrawal threshold from day 3 (RF; 22.02± 1.26, LF; 22.58± 1.33) to day 30 (RF; 17.00± 0.77, LF; 17.75± 1.06).

Intergroup comparison was done between all the groups and further t-test was applied between SCI and SCI+MF. There was significant increase in withdrawal threshold in forepaws at day 3, 10, and 30 in SCI group over sham and control, while there was no significant change in withdrawal threshold values on any of the days in hindpaw and forepaw when SCI+MF was compared with control and sham. Intergroup group comparison (t-test) between SCI and SCI+ MF group showed significant decrease in withdrawal threshold (g) in SCI+ MF (p value; RH 0.009, LH 0.005) group as compared to SCI in hindpaws on day 30. There was significant increase in withdrawal threshold (g) in SCI+ MF group as compared to SCI in forepaws at day 10 (p value; RF = 0.004, LF= 0.022) and day 30 (p value; RF = 0.002, LF= 0.002) (Figure 5)

**Figure 5:**
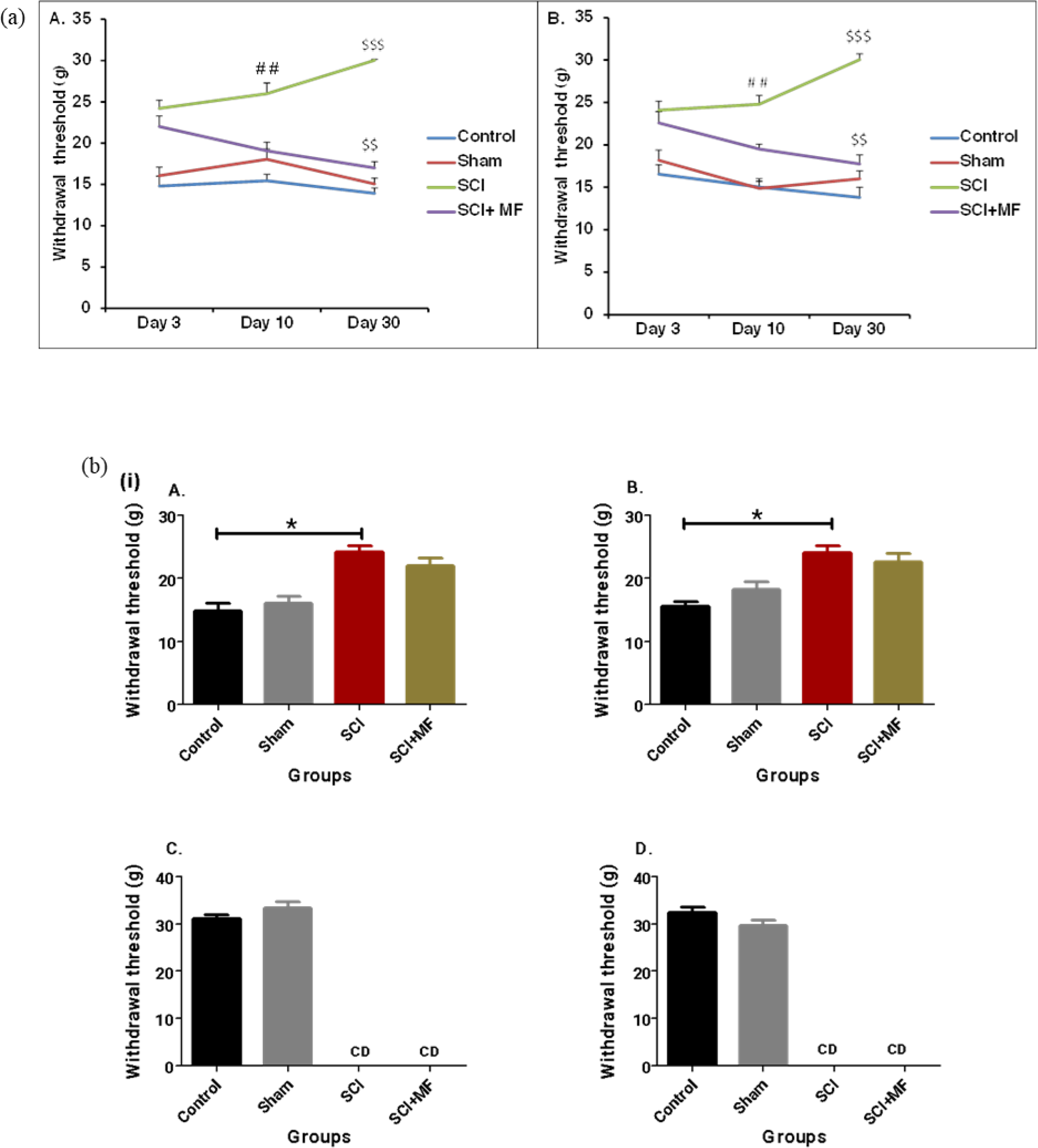
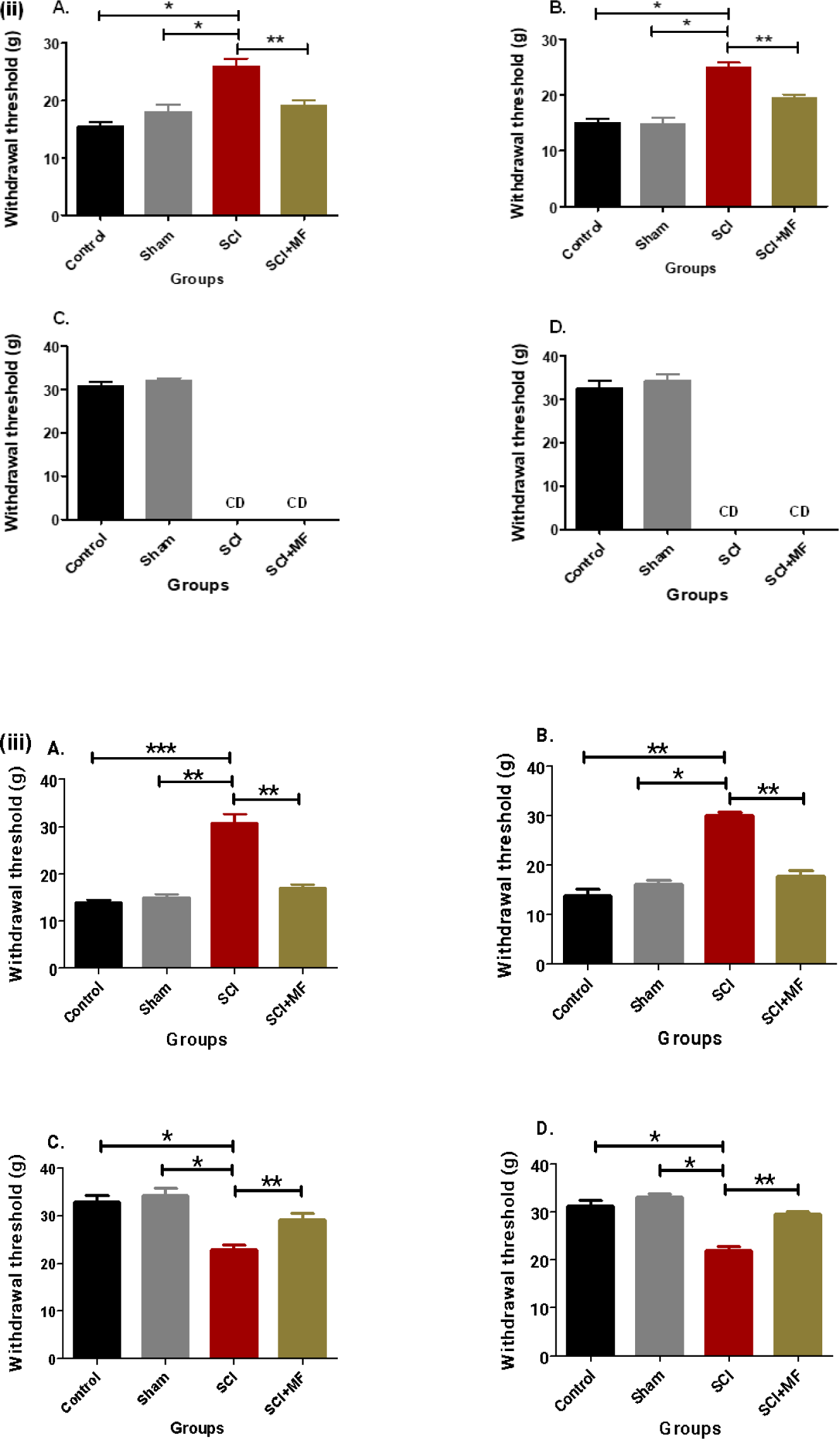
(a) Intragroup temporal comparison of withdrawal threshold (g) in right forepaw (A) and left forepaw (B); Intragroup temporal hindpaw graphs are not shown because the test could not be done (CD) on day 3 and 10. (b) Intergroup comparison of withdrawal threshold (g) at (i) Day 3, (ii) Day 10, (iii) Day 32 after MF exposure. Right forepaw (RF) (A); Left forepaw (LF) (B); right hindpaw (RH) (C); left hindpaw (LH) (D). * Indicates comparison between day 3 and day 10; ^#^ indicates comparison between day 10 and day 30; ^$^ indicates comparison between day 3 and day 30. Data expressed as Mean±SEM (n=6/ group). * p ≤ 0.05; ** p ≤ 0.01; *** p ≤ 0.001. CD= could not be done.

### Assessment of cortical electrical activity

Intragroup comparison was done to study the changes in power of different waves (δ, θ, α, and β) in both frontal and parietal lobe over the period of time using two way ANOVA. Intragroup temporal comparison of Frontal δpower (Fδp) and parietal δpower (Pδp), the time point (row factor; Frontal; p=0.111, F=2.33; df=2; Parietal; p=0.552,F=0.604, df=2) and δpower (column factor; F δp; p< 0.0001, F=17.62,df=3; P δp; p<0.0001; F=17.67; df=3) and the interaction between these two factors (Frontal; p= 0.018, F=2.95, df=6; Parietal; p= 0.902; F=0.356; df=6) showed significant increase in δpower from day 5 to 12 (p ≤ 0.01; from 3.66±0.89 to 10.26± 2.15 respectively) and significant decrease from day 12 to 32 in SCI group (p ≤ 0.01; from 10.26± 2.15 to 3.01± 0.62 respectively) in frontal lobe only, while in other groups there was no significant change in δpower over the period of time. Intragroup comparison of frontal θ power (Fθp) and parietal θ power (Pθp), the time point (row factor; Frontal; p=0.356, F=1.061; df=2; Parietal; p=0.843,F=0.171, df=2) and θ power (column factor; Fθp; p< 0.0001, F=13.87,df=3; Pθp; p< 0.0001; F=12.31; df=3) and the interaction between these two factors (Frontal; p=0.470, F=0.952, df=6; Parietal; p< 0.501; F=0.904; df=6) showed no significant change in θ power in any group over the time period in both the lobes. Intragroup comparison of frontal α power (Fαp) and parietal α power (Pαp), the time point (row factor; Frontal; p=0.896, F=0.109; df=2; Parietal; p=0.627,F=0.472, df=2) and α power (column factor; F αp; p< 0.0001, F=19.65,df=3; Pαp; p< 0.0001; F=14.31; df=3) and the interaction between these two factors (Frontal; p< 0.153, F=1.672, df=6; Parietal; p=0.605; F=0.761; df=6) showed no significant change in α power in any group over the time in both the lobes. Intragroup comparison of frontal βpower (Fβp) and parietal βpower (Pβp), the time point (row factor; Frontal; p=0.009, F=5.35; df=2; Parietal; p= 0.056, F=3.105, df=2) and βpower (column factor; Fβp; p=0.097, F=2.41,df=3; Pβp; p= 0.195; F=1.72; df=3) and the interaction between these two factors (Frontal; p=0.997, F=0.081, df=6; Parietal; p= 0.611; F=0.752; df=6) showed no significant change in βpower in any group over the time in both the lobes (Figure 6).

**Figure 6:**
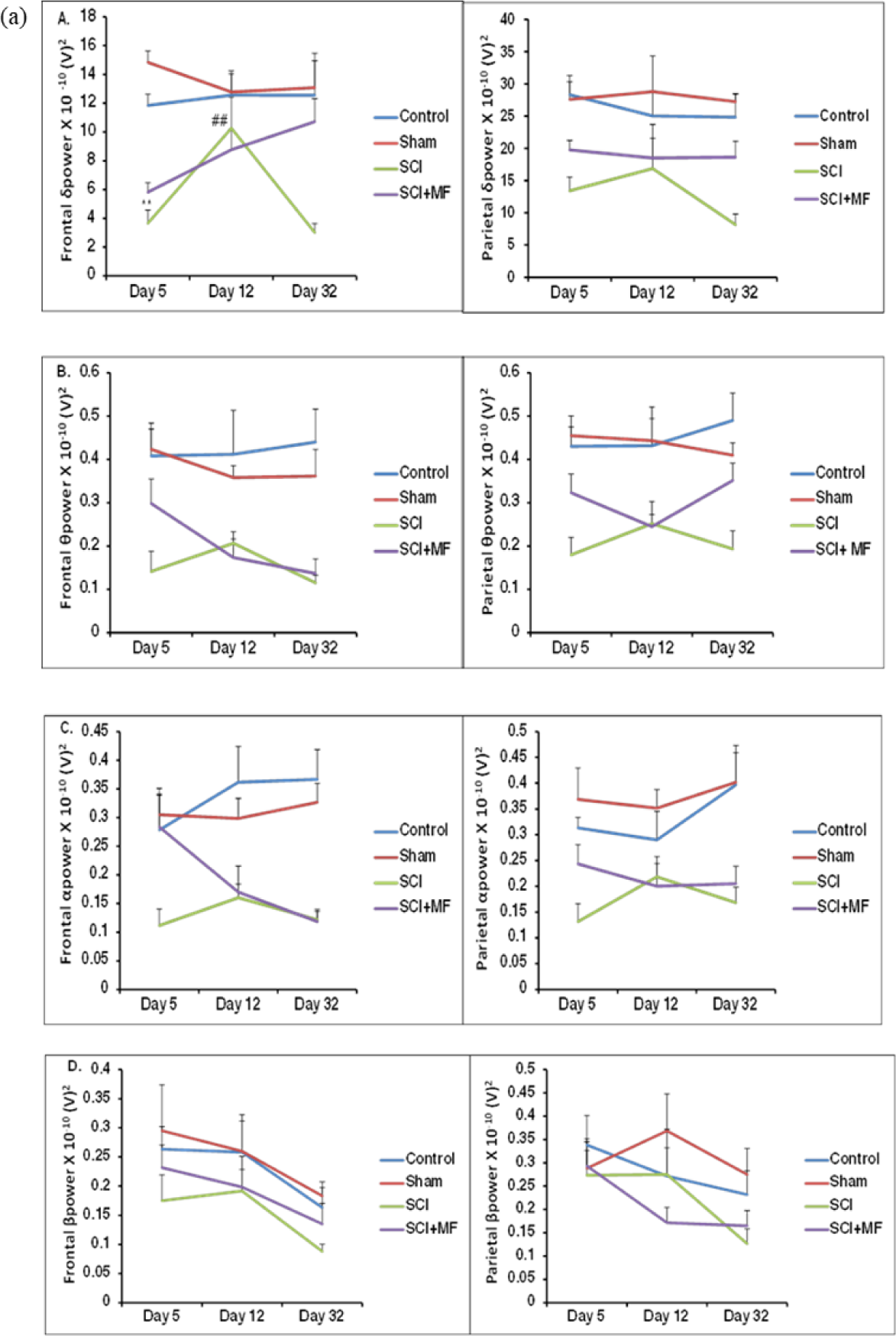
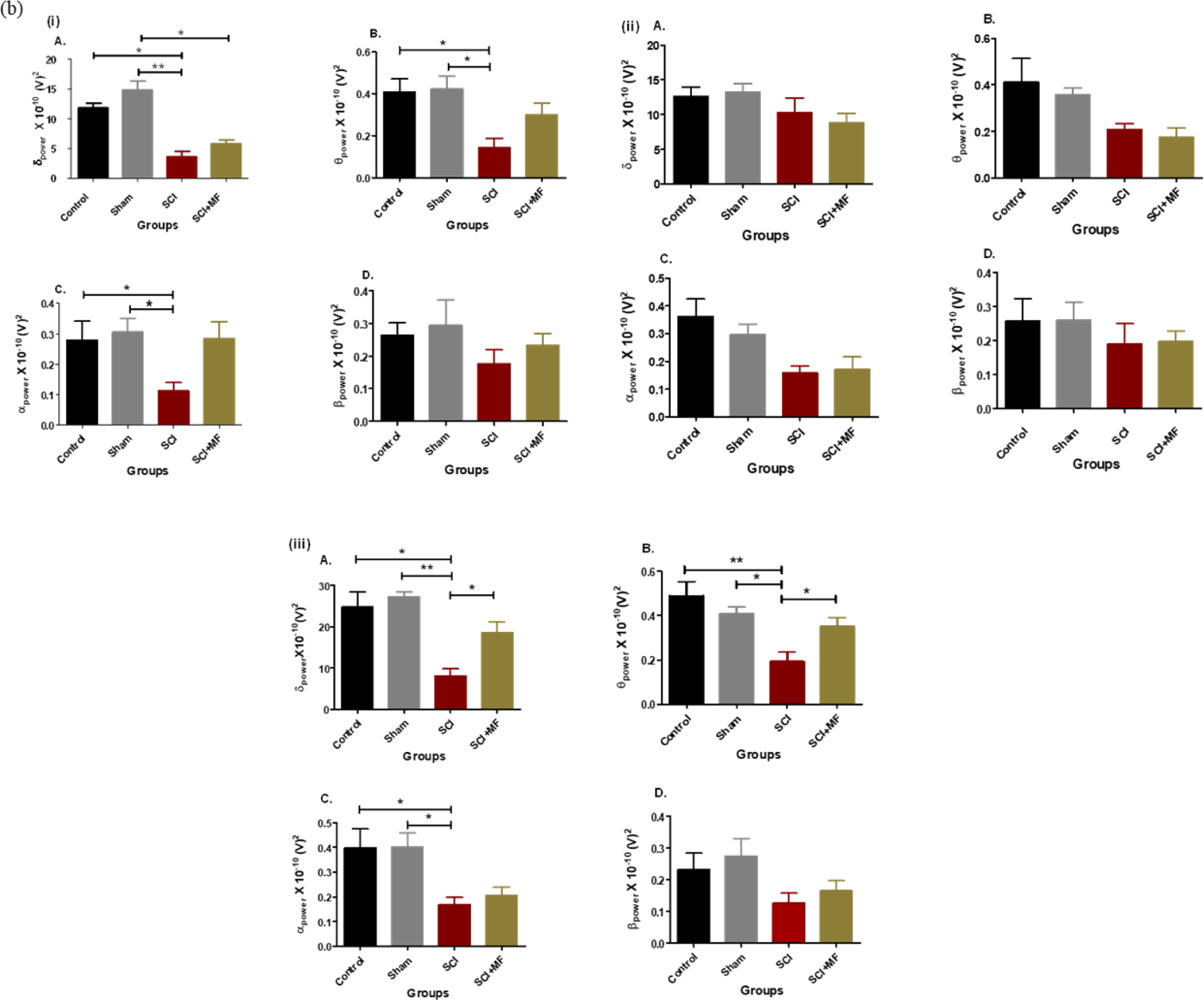
a) Intragroup temporal comparison of power (V)^2^ for different waves (δ, θ, α, and β) for frontal and parietal lobes. δpowerX10^-10^(V)^2^ for frontal and parietal lobes (A); θpowerX10^-10^(V)^2^ for frontal and parietal lobes (B); αpowerX10^-10^(V)^2^ for frontal and parietal lobes (C); βpowerX10^-10^(V)^2^for frontal and parietal lobes (D). (b) Intergroup comparison of power (V)^2^ for different waves (δ, θ, α, and β) in frontal lobe after MF exposure on (i) Day 5, (ii) Day 12, (iii) Day 32. δpowerX10^- 10^(V)^2^ (A); θpowerX10^-10^(V)^2^ (B); αpowerX10^-10^(V)^2^ (C), and βpowerX10^- 10^(V)^2^ (D). * Indicates comparison between day 5 and day 12; ^#^ indicates comparison between day 12 and day 32. Data expressed as Mean ±SEM (n=6/ group). * p ≤ 0.05; ** p ≤ 0.01; *** p≤ 0.001

Intergroup comparison was done between all the groups and further t-test was done between SCI and SCI+MF. Intergroup comparison showed decrease in δ, θ and α power in both frontal and parietal lobes in SCI group in comparison to control and sham at day 5 and day 32 but there was no change in β power in any of the days. Intergroup comparison between SCI and SCI+MF showed no significant change in any of the power (δ, θ, α, and β) after MF exposure in both the lobes at day 5 and 12. There was significant increase in δ power and θ power in frontal (δ; p= 0.002; θ; p= 0.873) and parietal (δ; p= 0.016; θ; p= 0.030) after MF exposure on day 32 (Figure 7).

**Figure 7:**
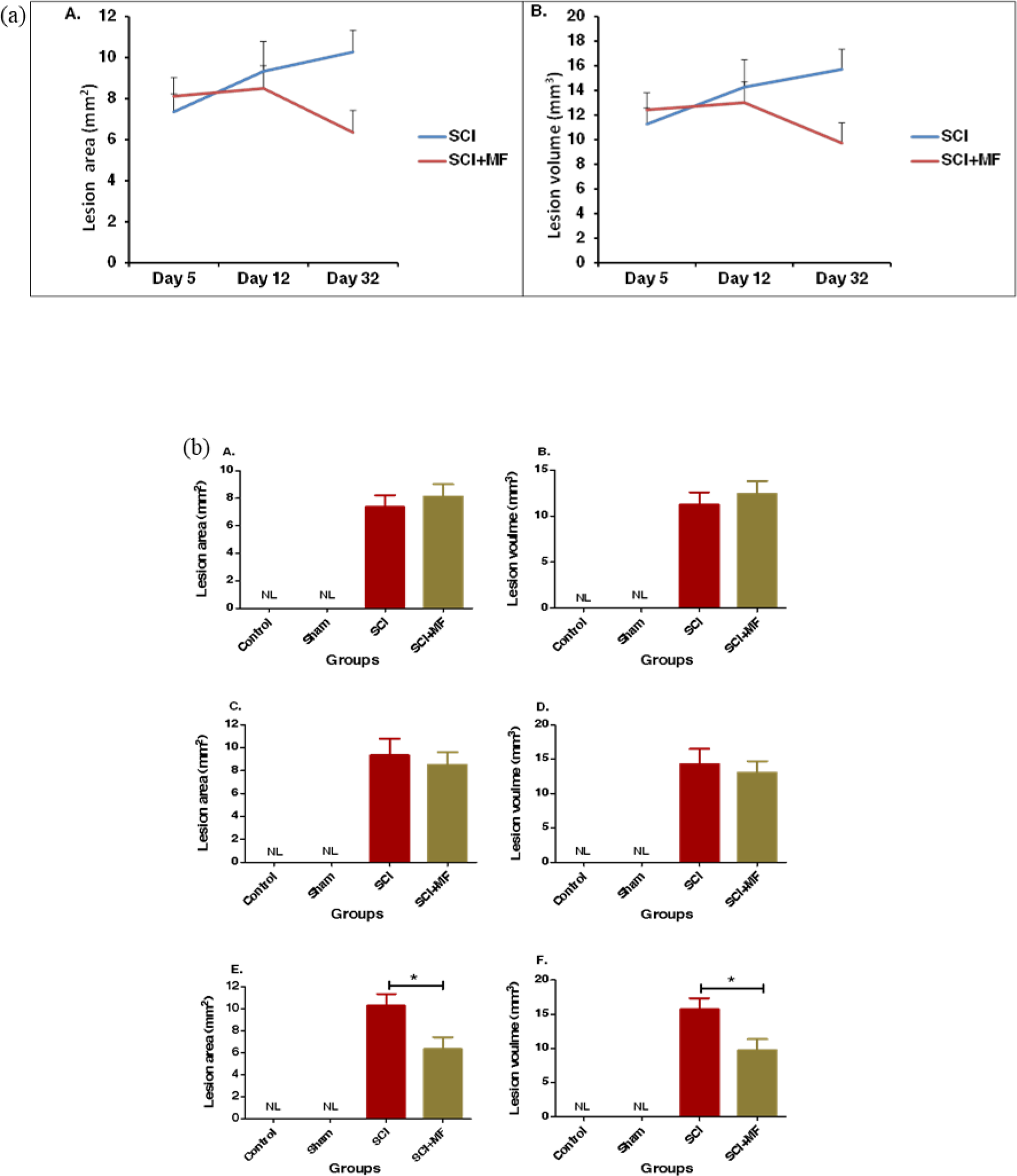
(a) Intragroup temporal comparison of lesion area (mm^2^) and lesion volume (mm^3^) in various groups. (b) Intergroup comparison of lesion area (mm) ^2^ and lesion volume (mm)^3^ after MF exposure. Day5 (A&B); day12 (C&D); day32 (E&F). Data expressed as mean± SEM (n=6/ group) * indicates comparison between SCI and SCI+MF groups; p≤ 0.05.

### Histological assessment

The histological assessments were performed at the end of the study period (day 5, 12 and 32) and the data obtained was compared in two ways: 1) Intragroup temporal comparison, and 2) Intergroup comparison.

### Assessment of lesion area and volume

On histological analysis no injury was observed in sham groups while in SCI and SCI+MF group, the staining of the damaged tissue (caudally and rostrally to epicenter of the injury) were indicative of the extent of the injury. The pathophysiological indication of the injury site in the isolated spinal cord tissues included penetration of inflammatory cells within white as well as gray matter, neuronal atrophy, hemorrhage, axonal damage, and cavities formation. Intragroup temporal comparison was done in SCI and SCI+ MF group to study the changes in the lesion area and volume over the period of time using two way ANOVA The time point (row factor; p= 0.542; F=0.631; df=2) and lesion area (column factor; p=0.201; F=1.87; df=1) and the interaction between these two factors (p= 0.101; F=2.58; df=2) found that changes in lesion area were not significant over the period of time. Intragroup temporal comparison of lesion area (mm^2^) in SCI group, showed no significant change from day 5 (7.36± 0.86 mm^2^) to day 32 (10.27±1.07 mm^2^) and in SCI+MF group also there was no significant change from day 5 (8.12± 0.91 mm^2^) to day 32 (6.35±1.08 mm^2^). Intragroup temporal comparison of lesion volume (mm^3^) in SCI, showed no significant change in lesion volume (mm^3^) from day 5 (11.26± 1.32 mm^3^) to day 32 (15.71± 1.63) and in SCI+MF group also there was no significant change from day 5 (12.42± 1.39mm^3^) to day 32 (9.72± 1.65mm^2^) (Figure7).

Intergroup comparison was done using t-test between SCI and SCI+MF, which showed significant decrease in lesion area (p=0.05) and volume (p=0.05) in SCI+MF in comparison to SCI group on day 32 but not on days 5 and 12, suggesting gradual recovery (Figure 7).

### Assessment of plasticity associated proteins (Nogo-A, BDNF)

Immunofluorescence was done for both Nogo-A and BDNF protein, images were taken at 40 X (Figure 8, 9), and counting was done using Image J software from different regions of the cortex.

**Figure 8:**
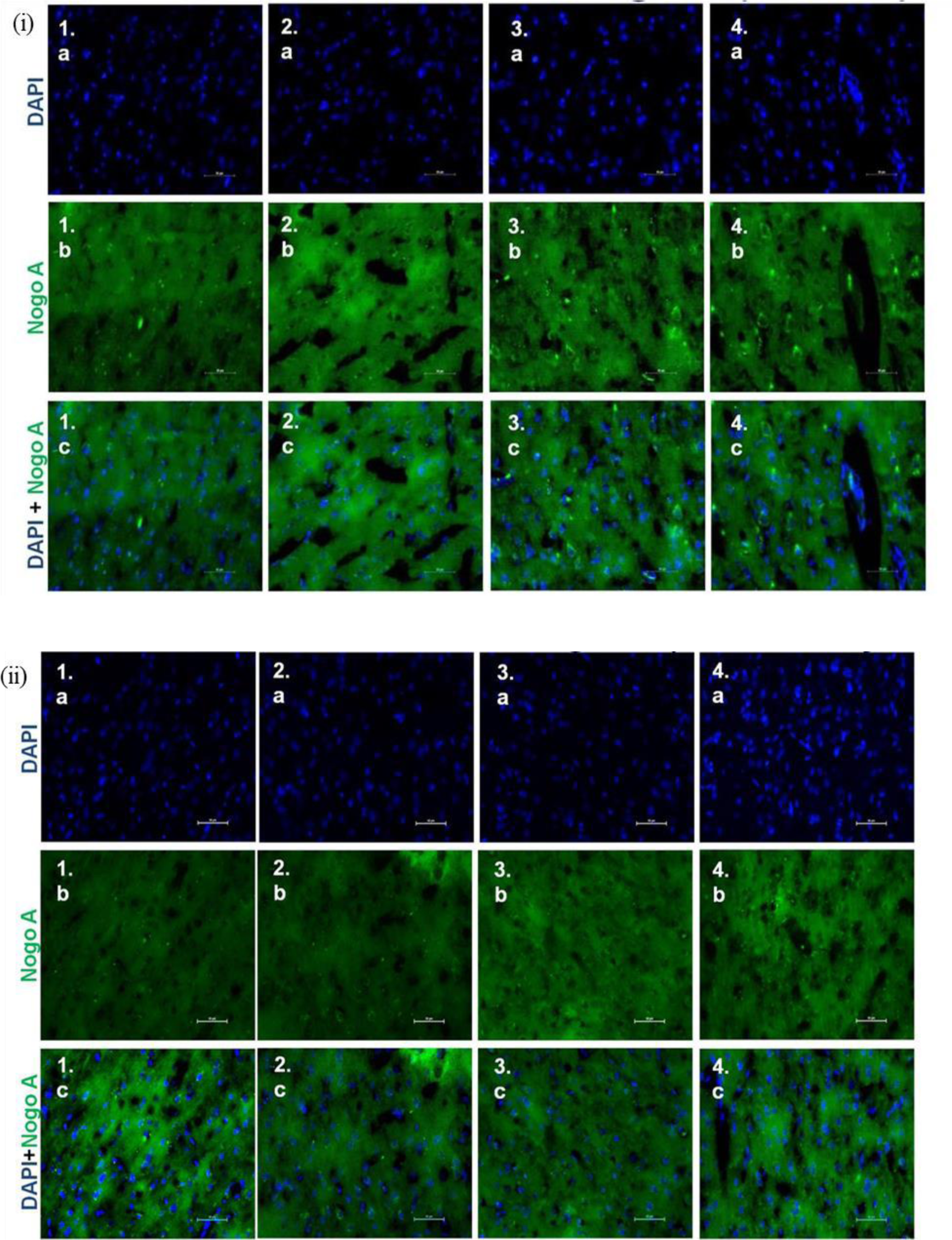
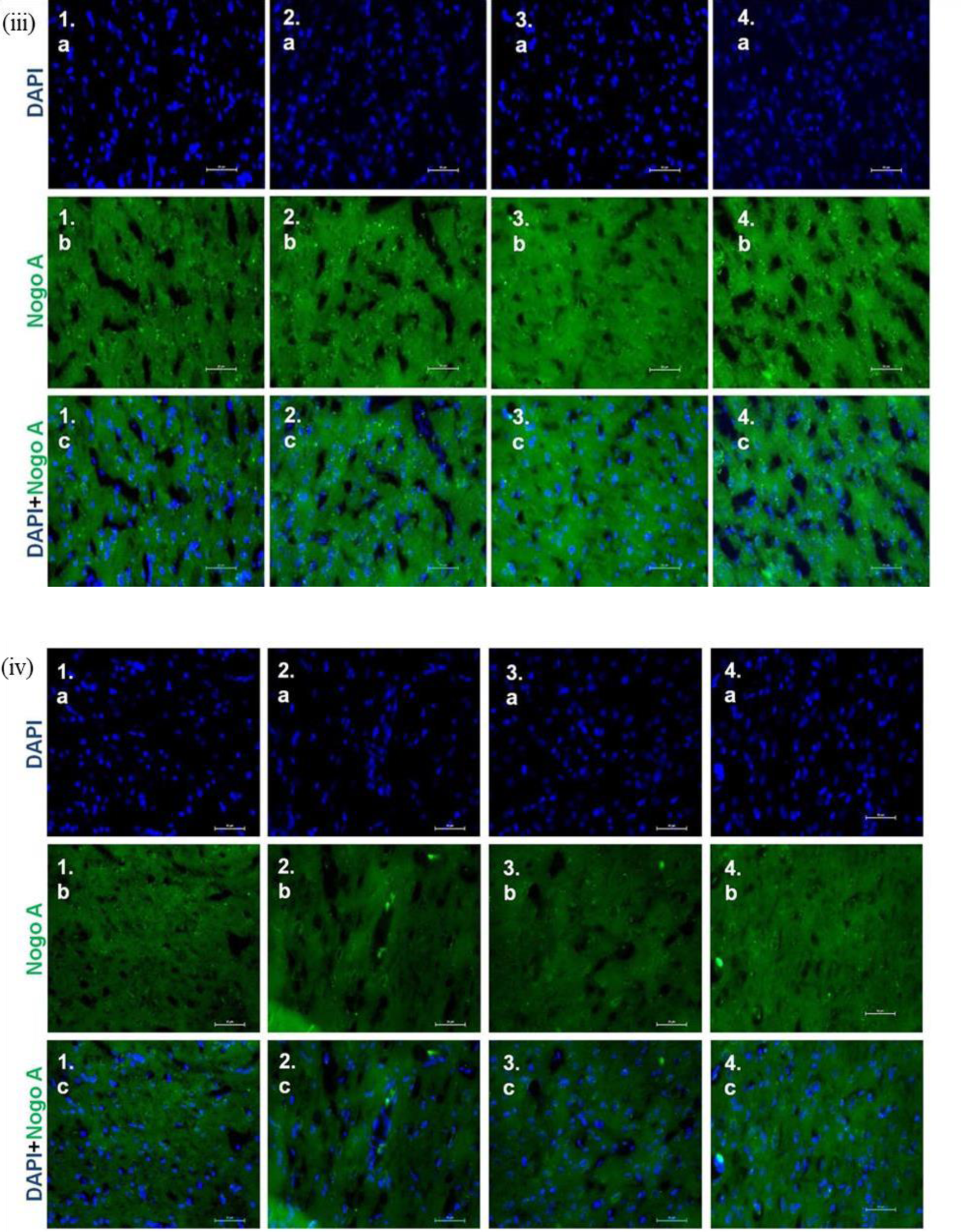
Subcellular localization of Nogo-A (40X objective lens) of M1 (primary motor area) (1a-c); M2 (secondary motor area) (2a-c); S1FL (somatosensory forelimb area) (3a-c); S1HL (somatosensory hindlimb area) (4a-c) in (i) Control group, (ii) Sham (Day 32), (iii) SCI (Day 32), (iv) SCI+MF (Day 32).

**Figure 9:**
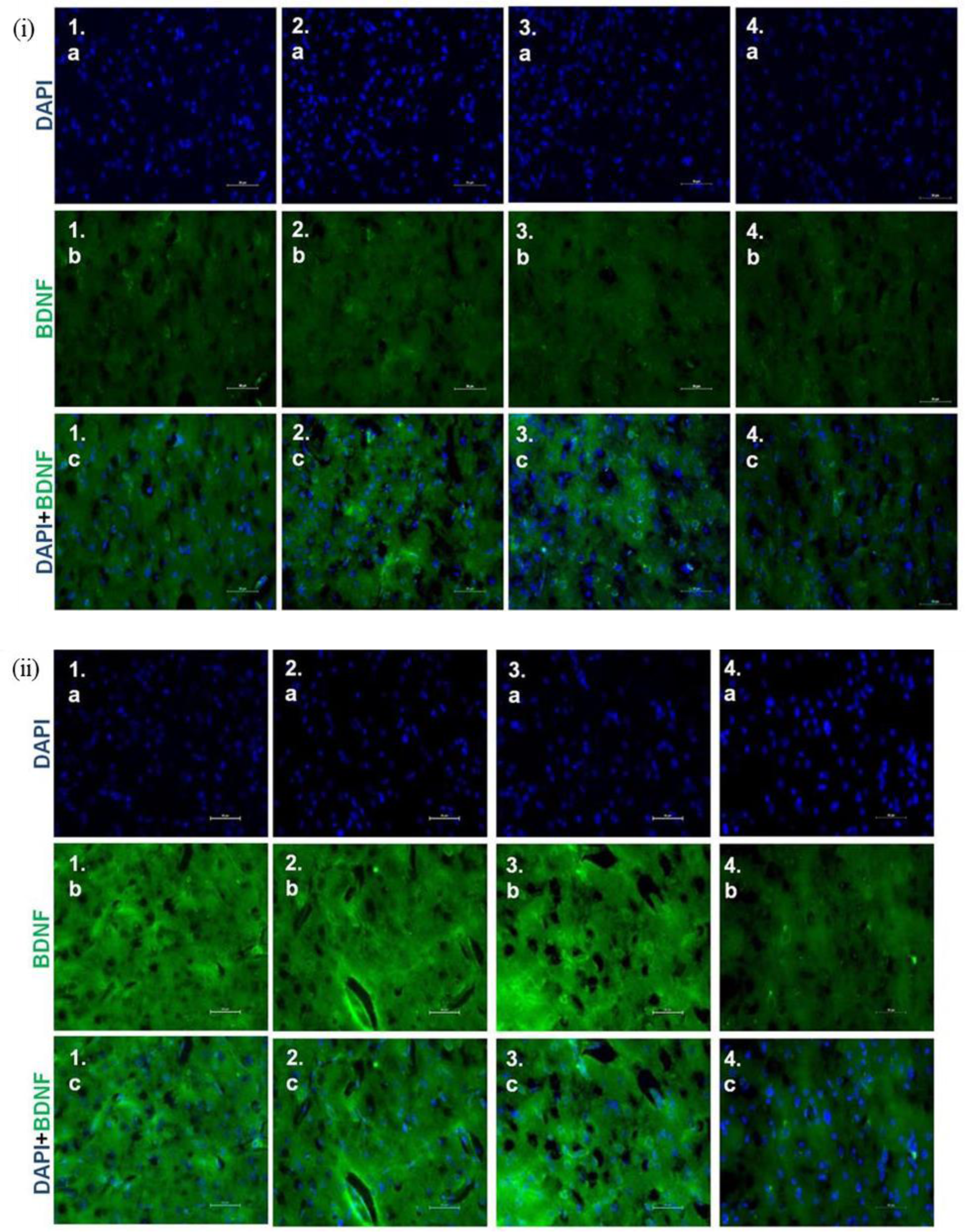
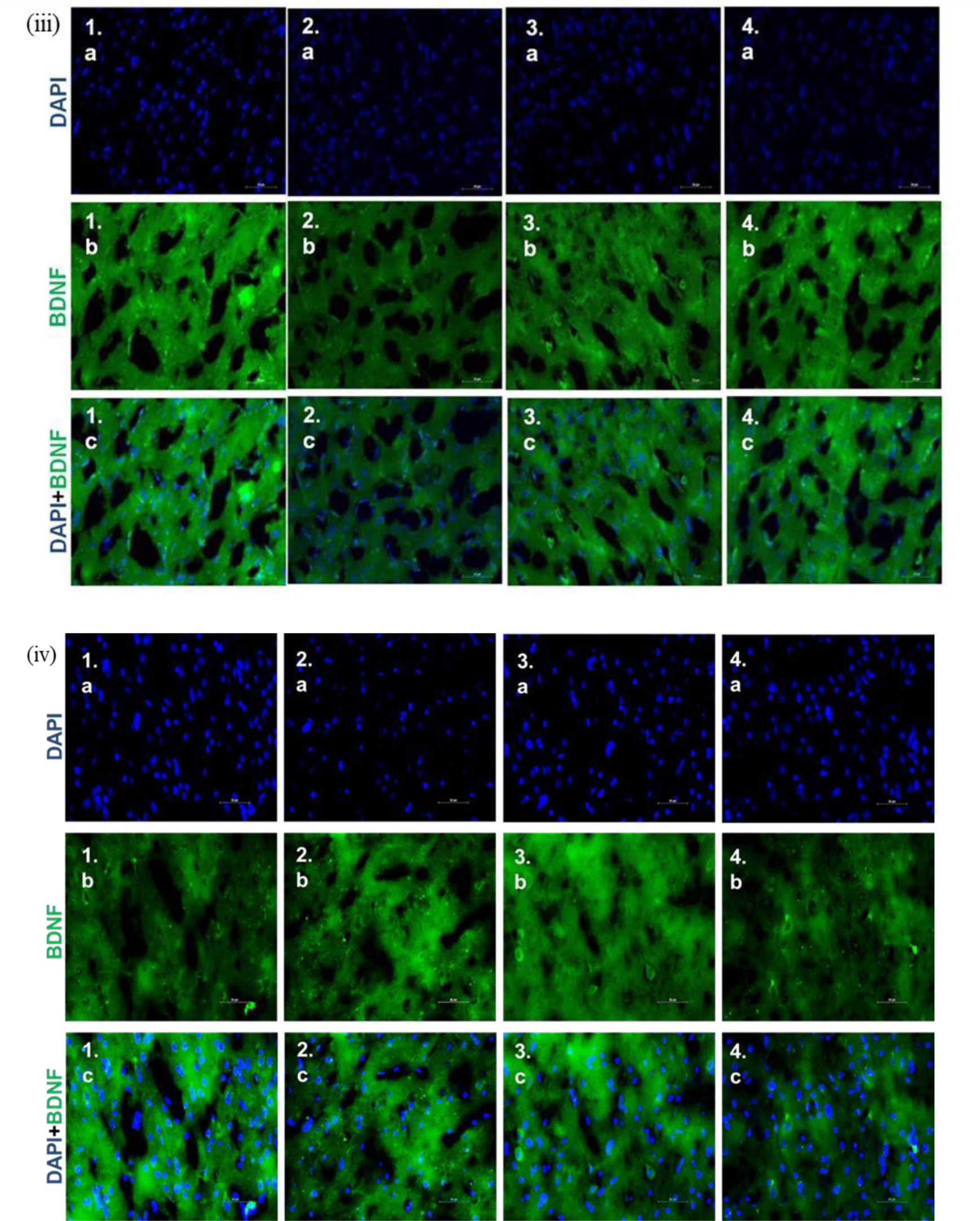
Subcellular localization of BDNF (40X objective lens) of M1 (primary motor area) (1a-c); M2 (secondary motor area) (2a-c); S1FL (somatosensory forelimb area) (3a-c); S1HL (somatosensory hindlimb area) (4a-c) in (i) Control group, (ii) Sham (Day 32), (iii) SCI (Day 32), (iv) SCI+MF (Day 32).

Two-way ANOVA was used to assess intragroup temporal comparison for the expression of Nogo-A and BDNF in all the groups. For Nogo-A protein, count/ field in right hemisphere in both SCI and SCI+MF group in M1 region, the time point (row factor; RH; p= 0.0007, F= 8.81, df=2; LH; p=0.0022,F= 7.59, df=2) and Nogo+ cells/ field (column factor; RH; p< 0.0001, F=17.73, df=3; LH; p< 0.0001, F= 15.78, df=3), and the interaction between these two factors (RH; p=.0007, F=4.94, df=6; LH; p=0.0003; F= 5.61; df=6), showed increase from day 5 to day 12 and then decreased on day 32. Intragroup temporal comparison of count/ field of Nogo-A protein in left hemisphere of M1as well as in both hemispheres of M2, S1FL and S1HL regions in both SCI and SCI+MF group also showed increase from day 5 to day 12 followed by a decrease on day 32, suggesting a biphasic response (Figure 10). There was no significant change in sham group.

**Figure 10:**
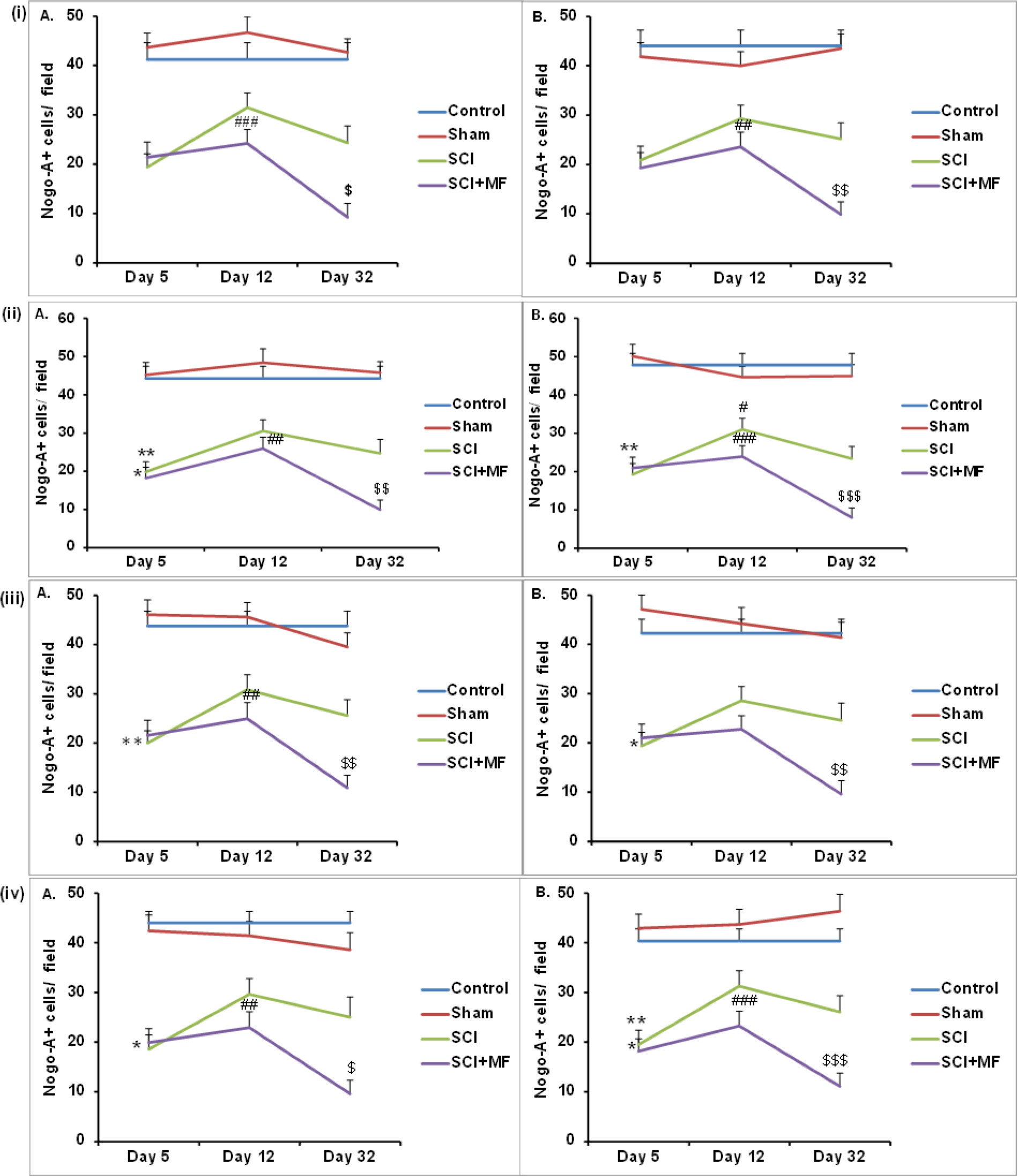
Intragroup temporal comparison of Nogo-A+ cells/ field of (i) M1 (primary Motor cortex), (ii) M2 (secondary motor cortex), (iii) S1FL (somatosensory, forelimb cortex), (iv) S1HL (somatosensory, hindlimb cortex) in right cortical hemisphere (A) and left cortical hemisphere (B). Data expressed as Mean±SEM (n=3/ group). * indicates comparison between day 5 and day 12; ^#^ indicates comparison between day 12 and day 32; ^$^ indicates comparison between day 5 and day 32. Field = 8730 μm^2^. ^*/#/$^ ≤0.05; ^**/##/$$^ ≤0.01, ^***/###/$$$^ ≤0.001.

Intergroup comparison for Nogo-A protein, showed a significant decrease (p≤0.05) in Nogo-A+ cell count/ field on day 5 only in comparison to control and sham group in all the regions of SCI. In SCI+MF, a significant decrease (p≤0.05) was observed at all the time points as compared to control and sham. A comparison between SCI vs SCI+ MF groups showed significant decrease (p≤0.05) in Nogo-A+ cell count/ field on day 32 but not at day 5 and 12. For BDNF protein, there was a significant increase (p≤0.05) in BDNF+ cell count/ field on day 5 in SCI in all the regions as compared to control and sham group. In SCI+MF, a significant increase was evident on day 12 (p≤0.05, only in M1, M2 and S1FL) and on day 32 (p≤0.01, in all the regions) as compared to control and sham. While SCI vs SCI+ MF group comparison showed significant increase in BDNF+ cell count/ field (p≤0.01) at day 32 in SCI+MF group but not on days 5 and 12 (Figure 11).

**Figure 11:**
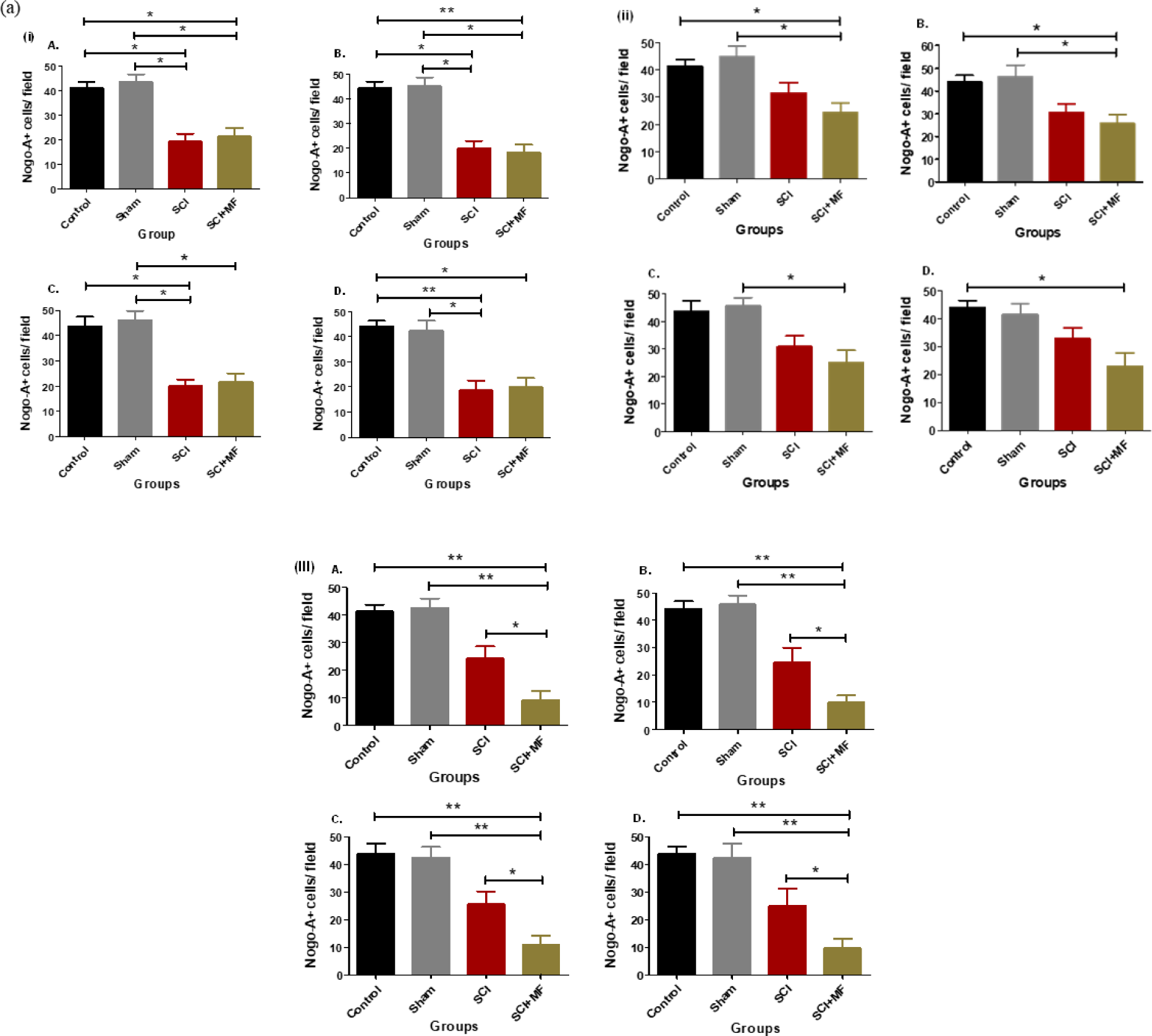
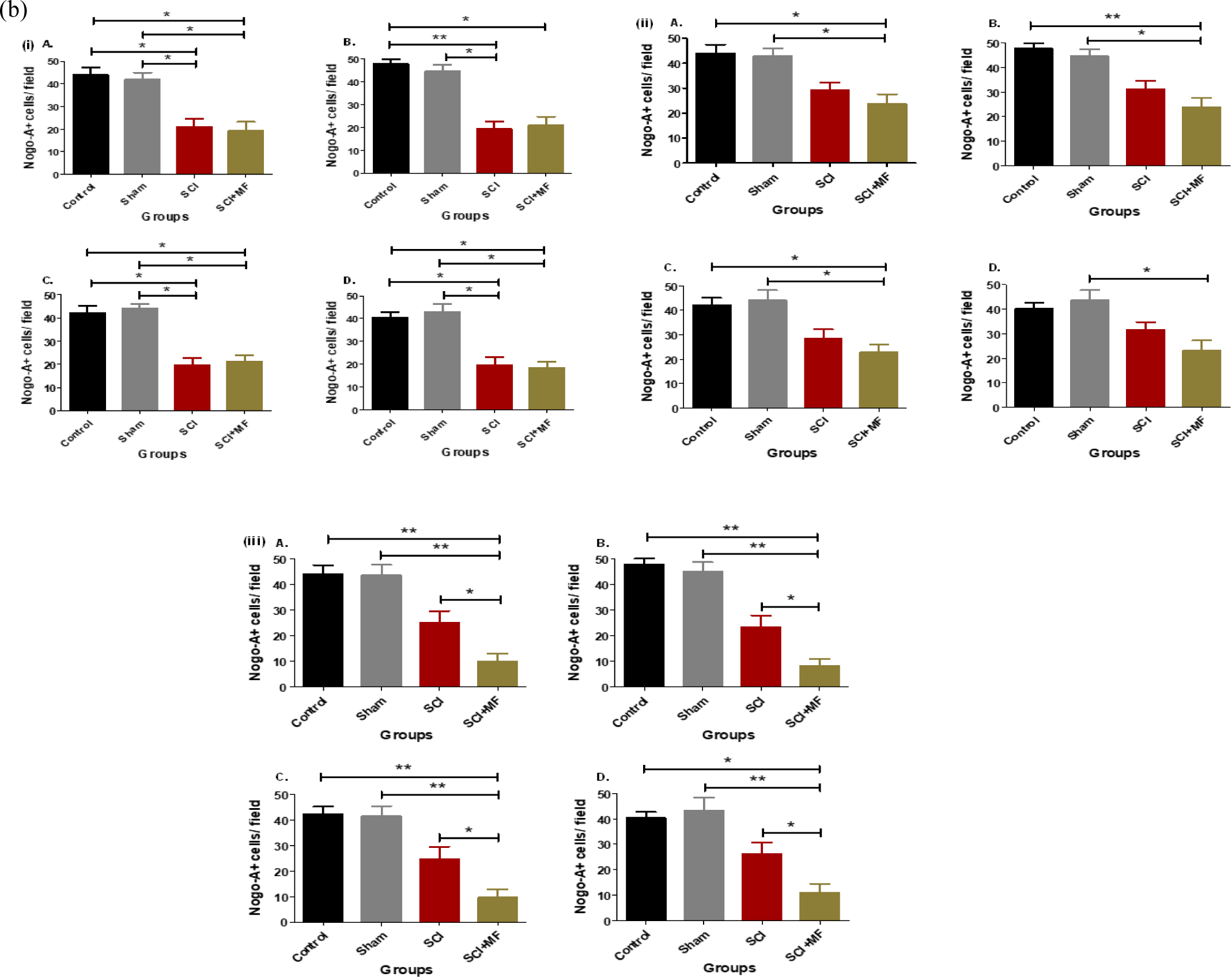
Intergroup comparison of Nogo-A+ cells/ field in (a) right and (b) left cortical hemisphere after MF exposure at (i) Day 5, (ii) Day 12 (iii) Day 32. M1= Primary Motor cortex (A); M2= Secondary Motor Cortex (B); S1FL= Primary somatosensory, forelimb (C); and S1HL= Primary somatosensory, hindlimb (D). Data expressed as Mean±SEM (n=3/ group). Field = 8730 μm^2^. * p ≤ 0.05; ** p ≤ 0.01; ***p ≤ 0.001

For BDNF protein in M1 region count/ field in right hemisphere in both SCI and SCI+MF groups, the time point (row factor; RH; p <0.0001, F= 74.74, df=2; LH; p<0.0001, F=129.1, df=2) and BDNF+ cells/ field (column factor; RH; p< 0.0001, F= 16.06, df=3; LH; p<0.0001, F=14.87, df=3), and the interaction between these two factors (RH; p<0.0001, F= 46.95, df=6; LH; p< 0.0001; F= 75.31; df=6), showed decrease from day 5 to day 12 and then increase on day 32. There was no significant change in sham group. Similar results were observed when intragroup temporal comparison of count/ field in left hemisphere of M1 and in both hemispheres of M2, S1FL and S1HL areas of both SCI and SCI+MF group were done, suggesting a biphasic response opposite in direction as observed for Nogo-A protein (Figure 12, 13).

**Figure 12:**
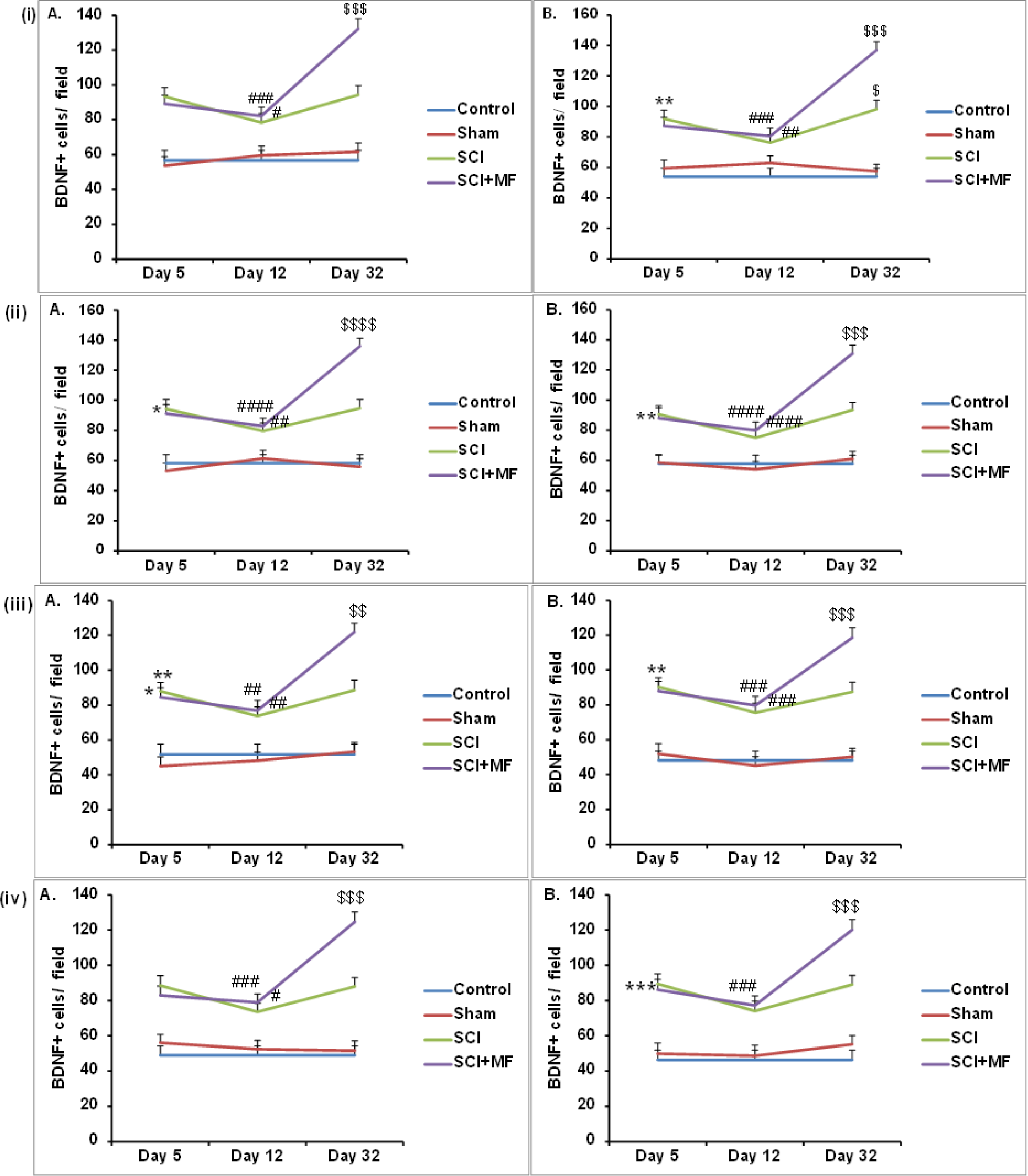
Intragroup temporal comparison of BDNF+ cells/ field of (i) M1 (primary Motor cortex), (ii) M2 (secondary motor cortex), (iii) S1FL (somatosensory, forelimb cortex), (iv) S1HL (somatosensory, hindlimb cortex) in right cortical hemisphere (A) and left cortical hemisphere (B). Data expressed as Mean±SEM (n=3/ group). * Indicates comparison between day 5 and day 12; ^#^ indicates comparison between day 12 and day 32; ^$^ indicates comparison between day 5 and day 32. Field = 8730 μm^2^. ^*/#/$^ ≤0.05; ^**/##/$$^ ≤0.01, ^***/###/$$$^ ≤0.001.

**Figure 13:**
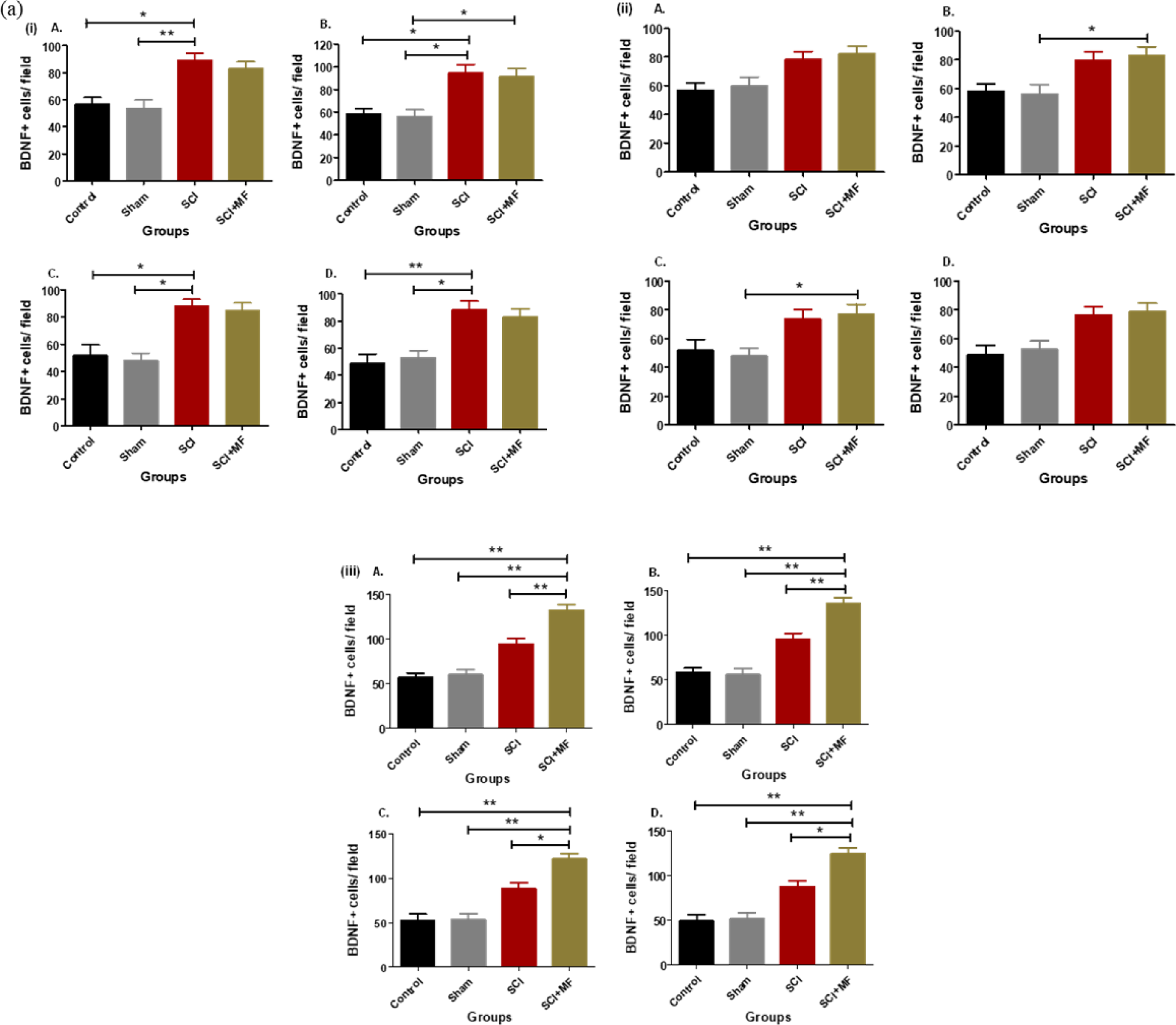
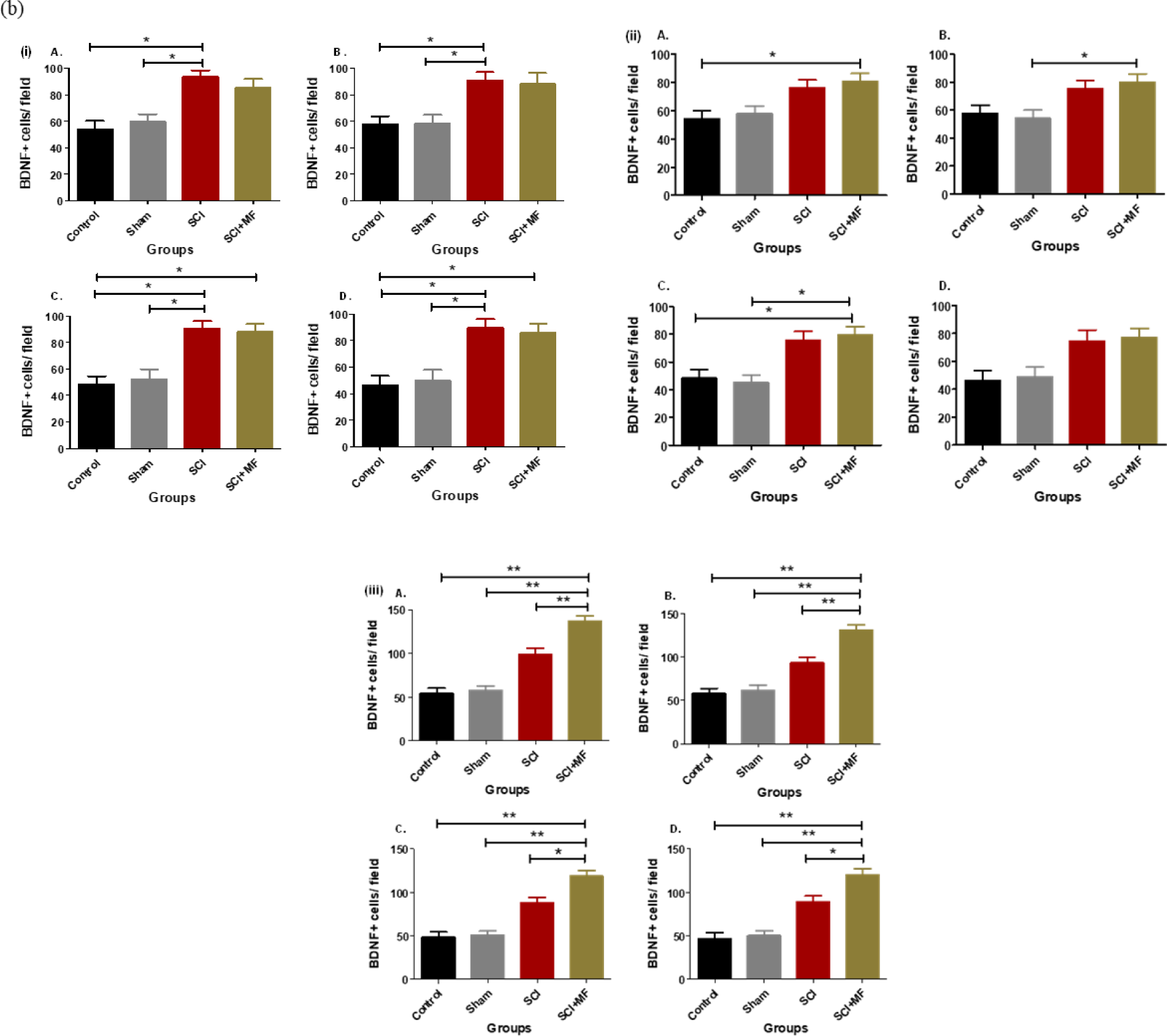
Intergroup comparison of BDNF+ cells/ field in (a) right (b) left cortical hemisphere after MF exposure at (i) Day 5, (ii) Day 12 (iii) Day 32. M1= Primary Motor cortex (A); M2= Secondary Motor Cortex (B); S1FL= Primary somatosensory cortex, forelimb (C); and S1HL= Primary somatosensory cortex, hindlimb (D). Data expressed as Mean±SEM (n=3/ group). Field = 8730 μm^2^. * p ≤ 0.05; ** p ≤ 0.01; ***p ≤ 0.001.

## Discussion

This study elucidates the effect of low intensity magnetic field (17.96µT, 50Hz) on the cortical reorganization after spinal cord injury in a complete transection rat model. In our study, magnetic field exposure ameliorated functional deficits, cortical electrical activity and modulated plasticity associated proteins in complete SCI rats. It is rationale to attribute the functional recovery after SCI to the neural plasticity as the structural repair after SCI takes much longer to emerge and requires morphology to manifest. SCI has received substantial attention among all the CNS injuries as a target for regenerative therapies but in context to neural plasticity, it is way less investigated. Complete SCI transection serves as a model for cortical plasticity as it completely silences the sensory input to one area of the motor and somatosensory cortex while leaving the inputs to neighboring area intact.

After complete transection SCI, we found that there was complete loss of sensory and motor functions below the level of lesion leading to paraplegic rats that moved by dragging the forelimbs. These rats developed bladder dysfunction, allodynia, and increased forelimb muscular strength. The histological assessment of isolated spinal cord tissue showed penetration of inflammatory cells in white and grey matter, neural atrophy, hemorrhage, axonal damage, and cavity formation. There was decrease in cortical electrical activity that showed temporal relationship paralleled with plasticity associated proteins Nogo-A and BDNF.

Akin to previous studies, MF exposure led to recovery in urinary bladder and locomotor function after 32 days exposure (Das et al., 2012., Pal et al., 2013., Bhattacharyya et al., 2020). Low intensity magnetic field can scavenge free radicals which decreases secondary injury, stimulates central pattern generators (CPGs) which aids in hindlimb locomotion (Porrier et al., 2004; Ahmed et al., 2011; Pal et al., 2013). MF exposure also restores muscle contractile properties and decrease muscular degeneration facilitating improvement in locomotion (Macias et al., 2000; Mert et al., 2006). The forelimb muscular strength of paraplegic rats increased from day 3 to day 30 after the complete transection injury. As the paraplegic rats were walking by dragging their forelimbs in the cage, due to continuous usage of the forelimb, the muscular strength increased over the period of time (Sandrow-Feinberg et al., 2009). However, MF restored the forelimb muscular strength on day 10 and 30 akin to sham due to improvement in hindlimb motor function as evident by BBB scoring. The improvement in hindlimb function after MF exposure can be attributed to the restoration of 5-HT. In our present study, we also found an increase in BDNF after MF exposure. 5-HT along with BDNF has been termed as a dynamic duo in synaptic plasticity, contributing to behavioral recovery in neurodegenerative disorders (Mattson et al., 2004). These two signals are known to act as co-regulatory molecules for each other with 5-HT stimulating expression of BDNF, while BDNF increases the survival and growth of the 5-HT neuron (Mamourias, L.A. et al., 1995; Meaney et al., 2000; Pei et al., 2003).

The clinical studies validate that 64-82% of SCI patients experience some form of neuropathic pain after the injury due to central sensitization. SCI leads to persistent elevation of neuronal activity, glial activation, and glutamatergic transmission which causes hyperexcitability of the dorsal horn neurons leading to maladaptive circuitry, aberrant processing of pain, and ultimately chronic neuropathic pain. These changes due to central sensitization led to allodynia and/or hyperalgesia (Hulsebosch et al., 2009; Latremoliere & Woolf, 2009, Watson et al., 2014). We did von Frey test for both hindpaws and forepaws to test if there is a development of allodynia below and above the level of injury after complete SCI. A significant decrease in withdrawal threshold in hindpaws of SCI rats at day 30 suggests development of allodynia. Similar results are also observed after spinal cord contusion (Yoon et al.,2004) and in spared nerve injury model (Richner et al.,2011) below the level of the lesion. After an injury to the spinal cord, glutamate is the major excitatory neurotransmitter that is released excessively leading to membrane depolarization via N-methyl-D aspartate (NMDA) receptors (Nagy et al., 2004). The co-release of peptidergic transmitters, such as substance P and CGRP, which are found in C-fibres along with glutamate, is responsible for a prolonged slow depolarization of the neuron and subsequent removal of the NMDA block, thus permitting wind-up to occur (Budai et al, 1996; Suzuki et al., 2003; Khasabov et al., 2005). In forepaws the threshold increased from day 3 to day 30 which could be due to extension of the cortical neuronal receptive fields to the hand area (Endo et al., 2007; Tandon et al., 2008; Aguilar et al., 2010; Yague et al., 2010; Kambi et al., 2014; Chand & Jain., 2015). This is the first report on recording of withdrawal thresholds using von Frey in forepaw after SCI. There are several studies which showed that extremely low frequency (ELF<300Hz) electromagnetic fields affect several neuronal activities, including memory, by influencing Ca2+ signaling enzymes in NMDA receptor dependent functions (Manikonda et al., 2007).A definitive link between ELF-MF exposure and analgesic processing in animals has established that exposure to a time-varying ELF-MF of 0.5 Hz (0.2–0.4 μT) abolishes the enhanced nocturnal analgesic response to morphine in mice (Kavaliers et al., 1984). The present study found an increase in withdrawal threshold of hindpaw after exposure of MF on day 30, suggesting an improvement in allodynia. MF exposure resulted in decrease in withdrawal thresholds which could be due to partial recovery in hindpaw locomotor function as discussed above.

SCI affects the electrical activity of the brain. We prominently found δ followed θ waves as it was done in anesthetized rat and α and β had low amplitude. It has been reported that decrease of δ and θ results from white matter disturbances in the CNS which could be due to multiple reasons including bilateral lesions, tumors, stroke, abscess, hematoma, and spinal cord injury (Fried et al., 2011). Previous study on compression SCI model showed that there was decrease in amplitude of δ, θ, α and β in anesthetized albino rat (Ram et al., 1987). Present study showed decrease in the power of the δ, θ, α and β on day 5, followed by a transitory increase on day 12, and again decrease on day 32. There was a biphasic trend in frontal and parietal lobe for delta wave though it was most significant in the frontal lobe. The other waves also showed a biphasic trend but the powers were so small to comment much on them. This transitory recovery in electrical activity on day 12 could be because of transitory release of glutamate from degenerative nerve endings which may have established thalamocortical pathways (Endo et al., 2007). MF exposure for 32 days resulted in improvement of the cortical electrical activity of the brain which can be attributed to the effect of magnetic field on primarily Ca^2+^ ion which causes changes in the rate of calmodulin dependent phosphorylation of myosin or due to K^+^ dependent biochemical pathways which are known to play a significant role in the regulation of the cellular activity. This is also due to the effect of MF on the cell membrane, causing depolarization and altering the permeability to ions (Shuvalova et al., 1991; Markov et al., 1993; Vorobyov et al., 1998; Mathie et al., 2003; Linda et al., 2009; Sallam 2012). Blank and Soo (1998) established that 50 Hz magnetic fields upregulate the activity of Na, K-ATPase and the cytochrome oxidase’s rate constant.

MF exposure for 32 days resulted in decrease in lesion area and volume suggesting decrease in secondary damage. In in vitro and in vivo EMF exposure activates axonal regeneration (Lai et all, 1993; Macias et al, 2000), reduces lesion volume, and protects against oxidative stress during secondary injury (Crowe et al, 2003; Ahmed et al, 2011). The electromagnetic field has been shown to have beneficial effects on wound healing and is known to reduce inflammation by decreasing COX-2, PGE-2 levels. It increases blood flow, modulates growth factor receptors, and also scavenges oxidative stressors like O2- (Pilla 2002). ELF-MF has also been shown to accelerate the repair process after inflammation (Patruno et al, 2017). Pulsed EMF has been recommended as therapy for post-operative pain and also for reduction of edema in osteoarthritis, and plastic surgeries (Finni et al, 2005; Thamsborg et al, 2005). Studies have also shown that PEMF can cause attenuation of scar formation after peripheral nerve injury in rats (Raji and Bowden 1983).

A biphasic pattern in expression of Nogo-A and BDNF was observed with a decrease in Nogo-A+ cells/ field on day 5, increase on day 12, and then again decrease on day 32, while BDNF+/ field cells increased on day 5, decreased on day 12 and then increased on day 32 in both SCI and SCI+MF group when compared with sham and control. The results are in tandem with the biphasic pattern of Nogo-A and BDNF established by Endo et al., (2007). The initial changes due to unanticipated and complete loss of sensory connection were replaced by short-term reactivation of the sensory inputs to the hindlimb cortex. This is due to the release of glutamate from degenerated sensory axons (Garrett and Thulin, 1975) that were projecting to the thalamus which lead to the transitory activation of the thalamocortical pathway to the hindlimb somatosensory cortex, followed by a permanent stoppage of this transitory input. A significant increase in BDNF and decrease in Nogo-A after MF exposure was observed only at day 32, suggesting effectiveness of MF when given for long durations. Previous studies have shown that magnetic field exposure promotes release of neurotrophic factors like BDNF in aberrant visual cortical circuitry (Makowieki et al., 2014), indicating upregulation of the BDNF as one of the major mechanism of action of the magnetic field induced plasticity (Gersner et al., 2011; Rodger et al., 2012). Transcranial magnetic stimulation has also been shown to increase the release of BDNF in patients with major depressive disorders (Xiao et al., 2018). Studies have shown that inducing BDNF pharmacologically or behaviorally caused reactivation of critical period-like plasticity in the mature visual system and lead to reorganization of the visual cortex (Mandolesi et al.,2005; Sale et al., 2007; Baroncelli et., 2010). Nogo-A signaling pathway is yet another major candidate for explaining the mechanism of cortical reorganization after spinal cord injury (Endo et al., 2007, 2009). Nogo-A, being a myelin-associated inhibitory factor is considered as a major therapeutic target in the last several years (Hulsebosch et al., 2002; Shwab et al., 2006; Akbik et al., 2012). Using an antibody against Nogo-A has been proved to be a promising therapy for promoting regeneration and plasticity in adult CNS (Schnell & Schwab, 1990; Z’Graggen et al. 1998; Blochinger et al., 2001; Emerick & Kartje 2004; Schwab 2004). Nogo-A antibodies given using osmotic minipumps intrathecally, increased regeneration of corticospinal neurons, behavioral improvement (Brosamole et al.,2000), and locomotor recovery (Merkler et al, 2001; Liebscher et al, 2005).

Brain stimulation paradigms that maximizes cortical reorganization after sensory and motor deafferentiation leads to increase in the activity of the intact cortex to compensate for the lost function (Penfield & Boldrey, 1937; Raineteau & Schwab, 2001; Freund et al., 2011; Romero et al., 2011; Bailey et al., 2015; Grover et al., 2015, van de rui & Grey, 2016) or put the overall cortex in more plastic state generically promoting cortical plasticity (Levy et al., 1990, Cohen et al., 1991, Streletz et al., 1995). It is therefore suggested that low intensity magnetic field (17.96 µT, 50Hz) given for 2h/ day for one month to complete spinal cord transected rats created more plastic state by regulating expression of Nogo-A and BDNF leading to functional reorganization that caused behavioral, morphological and electrical improvement. Magnetic field has been shown to induces electric currents to modulate cortical activity (Pascual-Leone, 2006), persistent excitability within cortical as well subcortical regions of the brain (Arias-Carrión, 2008) leading to behavioral modifications (Maeda et al., 2000; Valero-Cabré et al., 2005). It can produce a favorable microenvironment for neural repair by promoting neurotrophic factors, releasing neurotransmitters, axonal growth, regulate apoptosis and oxidative stress (Kumar et al., 2010; Das et al., 2012; Kumar et al., 2013; Pal et al., 2013). The *in vitro* studies have shown that it can raise intracellular calcium levels (Mathie et al., 2003; Matin L.Pall., 2013), modify cell metabolism, and also modify cytoskeleton (Berg, 1993). Animal studies also reported induction of long-term synaptic potentiation or depression, expression of receptors, plasticity-related genes modulation, overexpression of growth factors (Rodger et al., 2012). These systemic effects of magnetic field have a considerable impact on brain function as it causes changes in cell membrane properties, affects the expression of plasticity associated protein like stimulates BDNF and inhibits Nogo-A, thereby promoting cortical reorganization.

## Conclusion

The ability of magnetic field to induce electric field in the deeper underlying neural tissues creates conductive microenvironment that modulates the expression of plasticity associated proteins including BDNF and Nogo-A following SCI, leading to cortical reorganization favorably. Our study is unique as it unravels the biphasic effect of extremely low frequency magnetic field on sensory and motor cortical plasticity and electrical activity after complete spinal cord injury temporally until one month.

## Acknowledgments

We thank Dr. R.M. Pandey for the statistical analysis and Ms Priyanka Gupta for cresyl violet staining.

